# The worldwide invasion history of a pest ambrosia beetle inferred using population genomics

**DOI:** 10.1101/2023.01.25.525497

**Authors:** T. Urvois, C. Perrier, A. Roques, L. Sauné, C. Courtin, H. Kajimura, J. Hulcr, A.I. Cognato, M.-A. Auger-Rozenberg, C. Kerdelhué

**Affiliations:** INRAE, URZF, 45075 Orleans, France; UMR CBGP, INRAE, CIRAD, IRD, Institut Agro, Université Montpellier, Montpellier, France; Graduate School of Bioagricultural Sciences, Nagoya University, Nagoya, Japan; School of Forest, Fisheries, and Geomatics Sciences, University of Florida, Gainesville, FL, USA; Department of Entomology and Nematology, University of Florida, Gainesville, FL, USA; Department of Entomology, Michigan State University, East Lansing, MI, USA

**Keywords:** Bioinvasion, Invasion route, *Xylosandrus crassiusculus*, Genetic clusters, COI, RAD sequencing

## Abstract

*Xylosandrus crassiusculus*, a fungus-farming wood borer native to Southeastern Asia, is the most rapidly spreading invasive ambrosia species worldwide. Previous studies focusing on its genetic structure suggested the existence of cryptic genetic variation in this species. Yet, these studies used different genetic markers, focused on different geographical areas, and did not include Europe. Our first goal was to determine the worldwide genetic structure of this species based on both mitochondrial and genomic markers. Our second goal was to study *X. crassiusculus*’ invasion history on a global level and identify the origins of the invasion in Europe. We used a COI and RAD sequencing design to characterize 188 and 206 specimens worldwide, building the most comprehensive genetic dataset for any ambrosia beetle to date. The results were largely consistent between markers. Two differentiated genetic clusters were invasive, albeit in different regions of the world. The two markers were inconsistent only for a few specimens found exclusively in Japan. Mainland USA could have acted as a source for further expansion to Canada and Argentina through stepping-stone expansion and bridgehead events. We showed that Europe was only colonized by Cluster 2 through a complex invasion history including several arrivals from multiple origins in the native area, and possibly including bridgehead from the USA. Our results also suggested that Spain was colonized directly from Italy through intracontinental dispersion. It is unclear whether the mutually exclusive allopatric distribution of the two Clusters is due to neutral effects or due to different ecological requirements.

## Introduction

The number of biological invasions has increased in the last decades and is still increasing (Seebens et al. 2017, Sardain et al. 2019). Invasions are known to potentially have harmful effects on native biodiversity and ecosystems (Kenis et al. 2008, Simberloff et al. 2013) as well as on anthropized ecosystems (Paini et al. 2016) and human health (Jones and McDermott 2017). Invasion scenarios are diverse, ranging from a single introduction event (Hughes et al. 2017) to more complex histories implying multiple introductions and/or multiple sources (Javal et al. 2019, Tang et al. 2022). Moreover, invaded regions can act as sources for invasions to subsequent regions, a phenomenon called “bridgehead effect” (Lombaert et al. 2010, Bertelsmeier et al. 2021). Invasion history shapes the invasive populations’ genetic structure and diversity, which can affect the invasion dynamics. For example, invasions starting with a limited number of individuals and a low diversity of genotypes may result in significant genetic load and low evolutionary potential in the established populations (Schrieber and Lachmuth 2017). Identifying invasion routes and patterns of genetic structure and diversity are thus crucial to understand the demographic and genetic processes occurring in invasive populations and predict their progression, and to prevent further introductions via the identification of sources and points of entry. Studying worldwide genetic structure of invasive species can be considered as the most effective way to understand species’ invasion history (Estoup and Guillemaud 2010).

Studying population genetic structure of an invasive species can also reveal the existence of cryptic species or infraspecific genetic differentiation. Such information is crucial, as it can change the taxonomic scale at which the invasion should be considered and managed. For example, the *Euwallacea fornicatus* species complex was considered a single species when it was first described outside its native area, but is now considered to comprise seven species with overlapping morphologies (Smith et al. 2019). Studying its population genetic structure in Hawaii revealed the co-occurrence of two species of the complex (Rugman-Jones et al. 2020). Such findings can affect detection protocols and management decisions, for example, by helping find suitable natural enemies for biological control (Stouthamer et al. 2017). Information on infraspecific divergence is also essential, as different genetic lineages can have different biotic and abiotic preferences. Differentiated clades can have different potential distributions (Godefroid et al. 2015), potential impacts (e.g., if they preferentially attack different host plants) and origins.

The ambrosia beetles from the Xyleborini tribe are remarkable invaders, which can be explained by several biological characteristics (Hulcr and Stelinski 2017). They are minute species and they live inside galleries in their host plants. Hence they can be easily transported over long distances inside their hosts with international trade and escape sanitary inspections and treatments. They are xylomycetophagous (i.e. they feed on their symbiotic fungus rather than directly from the host plant tissues), which allows them to attack a broad range of host plant species. They are haplodiploid (i.e. haploid males hatch from non-fertilized eggs, while diploid females hatch from fertilized eggs), and they have a sib-mating reproduction, usually directly in their maternal galleries. This has several consequences. First, their genome is expected to be constantly purged from deleterious mutations, lowering the risk of inbreeding depression often observed in small invasive populations. Consistently, outbreeding depression but not inbreeding depression was previously documented in a species of ambrosia beetles (*Xylosandrus germanus*) (Peer and Taborsky 2005). Second, it eases mate finding even in very small populations, as during invasions. Lastly, haplodiploidy combined with their adult longevity allow single unmated females to establish a population, by mating with their male offsprings (produced from unfertilized eggs) to give a second generation comprising both haploid males and diploid females. Despite the damage caused by ambrosia beetles, the number of studies on their invasion history remains very limited. Kajtoch et al. (2022) reported less than 40 studies on saproxylic beetles’ phylogeography, and a particular lack of data for tropical and subtropical areas.

*Xylosandrus crassiusculus* is an ambrosia beetle originating from Southeastern Asia which is invasive worldwide. As opposed to most ambrosia beetles, *X. crassiusculus* attacks weakened and stressed trees in its invaded area, including fruit trees such as avocado (*Persea americana*) (Regupathy and Ayyasamy 2014), economically important crops such as cocoa (*Theobroma cacao*), coffee (*Coffea arabica*) and tea (*Camellia sinensis*), and ornamental trees such as *Cercis siliquastrum* (Kavčič and de Groot 2017). It invaded Africa more than a century ago (Hagedorn 1908, Schedl 1953), Pacific islands in 1950 (Samuelson 1981), North America in 1974 (Anderson 1974), South America in 2001 (Kirkendall 2018), and was detected in Europe recently, first in Italy in 2003 (Pennachio et al. 2003) and then in various European countries (France in 2014 (Roques et al. 2019), Spain in 2016 (Gallego et al. 2017), and Slovenia in 2017 (Kavčič 2018)). Several studies have already focused on *X. crassiusculus*’ genetic structure and phylogeography and suggested the existence of two genetically differentiated clusters (Dole et al. 2010, Landi et al. 2017, Storer et al. 2017, Nel et al. 2020). Ito and Kajimura (2009), focusing only on its native area in Japan, identified three distinct mitochondrial lineages. Despite these studies, the complete invasion picture remains unclear as the authors used different genetic markers, focused on different geographical areas, and did not include the European population.

This study aimed to characterize *Xylosandrus crassiusculus*’ genetic structure and decipher its worldwide invasion history. We used both a mitochondrial marker and pangenomic nuclear markers, as they can provide complementary information because they are differently inherited and do not evolve at the same rate (Toews and Brelsford 2012).

This approach will reconcile the results obtained in the previously published including individuals from Japanese clades and sub-clades (Ito & Kajimura (2009) and from the same regions as in Storer et al. (2017). Furthermore, we used the same mitochondrial fragment as in Landi et al. (2017) and Nel et al. (2020) so to include their data in our global analysis.

Our first goal was to compare the genetic structure based on the mitochondrial and nuclear markers and to determine whether they conformed to previous patterns, such as the two highly differentiated clusters identified worldwide by Storer et al. (2017) or the three mitochondrial lineages observed in Japan by Ito and Kajimura (2009). Our second goal was to study the invasion history of the species on a global-level, including the invasive populations in Europe. Specifically, we aimed to identify the origin(s) of the invasion in Europe, to determine whether it was invaded by several sources and whether bridgehead events occurred.

## Material & Methods

### Insect sampling

We assembled *Xylosandrus crassiusculus* females from 64 localities (Table 1, Supplementary Table 1), 31 localities in five countries in its native range and 33 localities in nine countries distributed on three continents in invaded ranges. While *X. crassiusculus* is often described as native to Eastern and Southeastern Asia (Ito and Kajimura 2009, Ranger et al. 2016, Gallego et al. 2017), the precise boundaries of its native range are unknown. We thus decided to consider all Asian localities as part of *X. crassiusculus* native area. The insects were either collected directly from the host tree, using traps baited with ethanol or more specific attractants (Roques et al., in prep), or obtained from collaborators. Whenever possible, individuals from the same location were selected as to minimize inter-individual relatedness within each location, by choosing different source trees or collections from different days. Individuals were stored in 96% ethanol and at -18°C until DNA extraction.

**Table 1:** Summary of the localities sampled and specimens used in the COI and RAD sequencing analyses. UFFE is short for University of Florida’s Forest Entomology Laboratory and uffeID represents the sample’s unique identifier in the UFFE collection database. PACA is short for the French region Provence-Alpes-Côtes d’Azur. The complete table featuring the latitude and the longitude of the different localities is available in Supplementary Figure 1.

| Sender (Genbank/accession number/uffeID) | Country | Locality | No. in COI analysis (haplotype) | Haplotype diversity | Nucleotide diversity | Mitochondrial cluster | No. In RAD analysis (major genomic group) |
| --- | --- | --- | --- | --- | --- | --- | --- |
| Genbank (KX685266.1) | Argentina | - | 1 (A02) | - | - | 2 | - |
| From Storer et al. 2017 | Cameroon | Limbe | - | - | - | 1 | - |
| Genbank (MF637141.1) | Canada | Point Pelee National Park | 1 (A02) | - | - | 2 | - |
| UFFE (19842) | China | Gushan | 1 (A04) | - | - | 2 | 1 (4) |
| UFFE (19847) | China | Nanan | 1 (E09) | - | - | 1 | 1 (9) |
| UFFE (19844) | China | Qishan | 1 (E02) | - | - | 1 | 1 (9) |
| UFFE (19848, 19851) | China | Shiwandashan | 2 (A11, E06) | 1 | 0.1093 | 1, 2 | 3 (4,9) |
| UFFE (19722) | China | Hainan | 5 (E01, E09) | 0.4 | 0.0113 | 1 | 6 (9) |
| Genbank (MN620075.1) | China | Hong Kong | 1 (E06) | - | - | 1 | - |
| UFFE (19878) | China | Kadoorie Farm | 1 (E06) | - | - | 1 | 1 (9) |
| UFFE (19877) | China | Tai Po Kau | 3 (E06) | 0 | 0 | 1 | 7 (9) |
| INRAE team | China | Wuhan | 4 (A11, A12) | 0.5 | 0.0044 | 2 | - |
| INRAE team | China | Xiangshan | 1 (E03) | - | - | 1 | 1 (9) |
| Genbank (MN620076.1) | China | Shanghai | 1 (A10) | - | - | 2 | - |
| Anthony Cognato | China | - | 2 (D06, D07) | 1 | 0.0018 | 1 | 5 (6) |
| From Storer et al. 2017 | China | Xishuangbanna | - | - | - | 1 | - |
| Jhuniar Morillo | Costa-Rica | Osa Peninsula | - | - | - | - | 1 (5) |
| UFFE (19874) | France | Carrefour de Gallion | 4 (D03) | 0 | 0 | 1 | 5 (7) |
| Genbank (KX055193.1, KX055195.1, KX055196.1) | France | Mont Marau | 3 (D02) | 0 | 0 | 1 | - |
| Genbank (KX055198.1) | France | Moorea Island | 1 (D02) | - | - | 1 | - |
| Genbank (KX055192.1) | France | Motu Tiahura | 1 (D02) | - | - | 1 | - |
| Genbank (KX055197.1) | France | Tahiti Island | 1 (D02) | - | - | 1 | - |
| INRAE team | France | Bayonne | 1 (A10) | - | - | 2 | - |
| INRAE team | France | Saint Jean de Luz | 2 (A03) | 0 | 0 | 2 | 2 (5) |
| INRAE team | France | Saint Maurice sur Adour | 2 (A03) | 0 | 0 | 2 | 5 (4) |
| INRAE team | France | Gignac | 1 (A10) | - | - | 2 | 1 (5) |
| INRAE team | France | Cap d'Ail | 5 (A10) | 0 | 0 | 2 | 3 (5) |
| INRAE team | France | Cap Ferrat | 1 (A10) | - | - | 2 | 2 (5) |
| INRAE team | France | Menton | 1 (A10) | - | - | 2 | 1 (5) |
| INRAE team | France | Mont Boron | 1 (A10) | - | - | 2 | 4 (5) |
| Genbank (MN182983.1, MN183016.1) | France | Nice | 2 (A10) | 0 | 0 | 2 | - |
| INRAE team | France | Nice Albert 1er | 2 (A10) | 0 | 0 | 2 | 2 (5) |
| INRAE team | France | Nice Vallée du Var | 1 (A10) | - | - | 2 | 1 (4) |
| INRAE team | France | Sainte-Marguerite | 3 (A10) | 0 | 0 | 2 | 3 (5) |
| INRAE team | France | Villa Thuret | 7 (A10) | 0 | 0 | 2 | 5 (5) |
| From Storer et al. 2017 | Ghana | Ankasa | - | - | - | 1 | - |
| From Storer et al. 2017 | Honduras | - | - | - | - | 2 | - |
| Genbank (MN620071.1) | India | Dehra Dun | 1 (D01) | - | - | 1 | - |
| UFFE (7942) | Indonesia | Bangunrejo | 2 (D02) | 0 | 0 | 1 | 5 (7) |
| UFFE (7954) | Indonesia | Blawan | 2 (D02) | 0 | 0 | 1 | 5 (7) |
| INRAE team | Italy | Parco Monti Aurunci | 3 (A02) | 0 | 0 | 2 | 5 (4) |
| INRAE team | Italy | Parco Riviera di Ulisse | 2 (A02) | 0 | 0 | 2 | 2 (4) |
| INRAE team | Italy | Sabaudia | 3 (A02) | 0 | 0 | 2 | 5 (3) |
| Hisashi Kajimura | Japan | Amami | 2 (A02, C04) | 1 | 0.0705 | 2 | 1 (2) |
| From Ito & Kajimura 2009 | Japan | Amami | - | - | - | 2 | - |
| Hisashi Kajimura | Japan | Okinawa, Naha | 3 (A07, C05, C06) | 1 | 0.0470 | 2 | 3 (2,4) |
| From Ito & Kajimura 2009 | Japan | Naha | - | - | - | 2 | - |
| From Storer et al. 2017 | Japan | Aichi | - | - | - | 2 | - |
| From Ito & Kajimura 2009 | Japan | Nagano, Chisagata-gun | - | - | - | 2 | - |
| Hisashi Kajimura | Japan | Nagano, Chisagata-gun | 2 (B06, B07) | 1 | 0.0018 | 2 | 2 (1) |
| From Ito & Kajimura 2009 | Japan | Nagano, Shimominochi-gun | - | - | - | 2 | - |
| From Ito & Kajimura 2009 | Japan | Nagano, Shiojiri | - | - | - | 2 | - |
| Hisashi Kajimura | Japan | Tottori | 2 (A02) | 0 | 0 | 2 | 2 (3) |
| From Ito & Kajimura 2009 | Japan | Tottori | - | - | - | 2 | - |
| Hisashi Kajimura | Japan | Hiroshima, Miyoshi | 2 (A02, B02) | 1 | 0.0705 | 2 | 2 (1,3) |
| From Ito & Kajimura 2009 | Japan | Hiroshima, Miyoshi | - | - | - | 2 | - |
| Hisashi Kajimura | Japan | Sapporo | 2 (B03, B05) | 1 | 0.0053 | 2 | 2 (1) |
| From Ito & Kajimura 2009 | Japan | Sapporo | - | - | - | 2 | - |
| From Ito & Kajimura 2009 | Japan | Toyama, Nakashinkawa-gun | - | - | - | 2 | - |
| From Ito & Kajimura 2009 | Japan | Saitama, Chichibu | - | - | - | 2 | - |
| From Ito & Kajimura 2009 | Japan | Chiba, Kamogawa | - | - | - | 2 | - |
| From Ito & Kajimura 2009 | Japan | Kyoto, Maizuru | - | - | - | 2 | - |
| From Ito & Kajimura 2009 | Japan | Kyoto, Nantan | - | - | - | 2 | - |
| From Ito & Kajimura 2009 | Japan | Mie, Isshi-gun | - | - | - | 2 | - |
| From Ito & Kajimura 2009 | Japan | Wakayama, Tanabe | - | - | - | 2 | - |
| From Ito & Kajimura 2009 | Japan | Miyakazi, Kobayashi | - | - | - | 2 | - |
| Hisashi Kajimura | Japan | Miyakazi, Kobayashi | - | - | - | - | 1 (1) |
| Genbank (MN620077.1) | Japan | Okinawa | 1 (D02) | - | - | 1 | - |
| Hisashi Kajimura | Japan | Koshi, Hata-gun | 2 (B01, B02) | 1 | 0.0035 | 2 | 3 (1) |
| From Ito & Kajimura 2009 | Japan | Koshi, Hata-gun | - | - | - | 2 | - |
| From Storer et al. 2017 | Japan | Iriomote | - | - | - | 2 | - |
| Hisashi Kajimura | Japan | Ishigaki | 3 (C01, C02, C03) | 1 | 0.0035 | 2 | 3 (2) |
| From Ito & Kajimura 2009 | Japan | Ishigaki | - | - | - | 2 | - |
| From Ito & Kajimura 2009 | Japan | Iwate, Iwate-gun | - | - | - | 2 | - |
| Hisashi Kajimura | Japan | Yamagata, Tsuruoka | 3 (B04, B05) | 0.667 | 0.0012 | 2 | 2 (1) |
| From Ito & Kajimura 2009 | Japan | Yamagata, Tsuruoka | - | - | - | 2 | - |
| Hisashi Kajimura | Japan | Shizuoka | 2 (A02) | 0 | 0 | 2 | 2 (3) |
| From Ito & Kajimura 2009 | Japan | Shizuoka | - | - | - | 2 | - |
| From Ito & Kajimura 2009 | Japan | Aichi, Toyota | - | - | - | 2 | - |
| Genbank (GU808709.1) | Madagascar | - | 1 (D02) | - | - | 1 | - |
| From Storer et al. 2017 | Madagascar | Ranomafana | - | - | - | 1 | - |
| Ben Boyd | New Zealand | Auckland | 4 (A02) | 0 | 0 | 2 | 5 (3) |
| Genbank (KX035196.1) | Panama | - | 2 (D04) | 0 | 0 | 1 | - |
| Anthony Cognato | Panama | Barro Colorado Island | 2 (D04) | 0 | 0 | 1 | 5 (5) |
| UFFE (19876) | Papua New | Kugofanka | 4 (A02, D02) | 0.5 | 0.0547 | 1, 2 | 5 (3,8) |
|  | Guinea |  |  |  |  |  |  |
| From Storer et al. 2017 | Papua New Guinea | Madang | - | - | - | 1 | - |
| UFFE (19863, 19864, 19865, 19866, 19867) | Papua New Guinea | Ohu | 2 (D10) | 0 | 0 | 1 | 5 (7) |
| Andreja Kavčič | Slovenia | Podsabotin | 16 (A09, A10) | 0.233 | 0.0025 | 2 | 5 (4) |
| From Nel et al. 2020 | South Africa | - | - | - | - | 1 | - |
| From Nel et al. 2020 | South Africa | Tzaneen | 3 (A06) | 0 | 0 | 2 | - |
| From Nel et al. 2020 | South Africa | - | 1 (D04) | - | - | 1 | - |
| Diego Gallego | Spain | El Tello | 12 (A02) | 0 | 0 | 2 | 5 (3) |
| Diego Gallego | Spain | Naquera | 2 (A02) | 0 | 0 | 2 | 5 (3) |
| UFFE (8515, 8520) | Taiwan | Dayueshan | 2 (E03) | 0 | 0 | 1 | 3 (7,9) |
| From Storer et al. 2017 | Taiwan | Dali | - | - | - | 1, 2 | - |
| Genbank (MN620074.1) | Taiwan | - | 1 (E10) | - | - | 1 | - |
| Genbank (GU808708.1) | Thailand | - | 1 (D05) | - | - | 1 | - |
| From Storer et al. 2017 | Thailand | Doi Pui | - | - | - | 1 | - |
| Genbank (BBCCA4264-12) | USA | Toad Suck Park | 1 (A02) | - | - | 2 | - |
| Genbank (BBCCA4147-12C) | USA | Collier Seminole State Park | 1 (A06) | - | - | 2 | - |
| Jared Bernard | USA | Manoa Valley | 4 (A01, A02, D02) | 0.833 | 0.0738 | 1, 2 | 5 (3,7) |
| Jared Bernard | USA | Moloka'a Bay | 4 (A02, A10) | 0.667 | 0.0129 | 2 | 4 (7) |
| Jared Bernard | USA | Poamoho Ridge | 3 (A05, D02) | 0.667 | 0.0670 | 1, 2 | 2 (4,7) |
| Anthony Cognato | USA | Vulcano Nat. Park | 2 (A02) | 0 | 0 | 2 | 5 (7) |
| Genbank (GU808710.1) | USA | - | 1 (A02) | - | - | 2 | - |
| Jhuniar Morillo | USA | Long Island | 2 (A02) | 0 | 0 | 2 | 2 (3) |
| Anthony Cognato | USA | Smithtown | 2 (A02) | 0 | 0 | 2 | 2 (3) |
| Genbank (GU808711.1) | USA | - | 1 (A02) | - | - | 2 | - |
| UFFE (20193) | USA | Cherokee | 2 (A02) | 0 | 0 | 2 | 5 (3) |
| Steve Frank | USA | Raleigh | 4 (A02, A07) | 0.5 | 0.0071 | 2 | 3 (3,4) |
| UFFE (17525, 17526) | USA | Leone | 8 (D02, D03) | 0.25 | 0.0004 | 1 | 5 (7) |
| UFFE (20230) | USA | Cosby | 2 (A02) | 0 | 0 | 2 | 5 (3) |
| Genbank (GMGSC323-12) | USA | Great Smoky Mountains Nat. Park | 1 (A02) | - | - | 2 | - |
| Anthony Cognato | USA | Tomball | 2 (A02, A08) | 0 | 0 | 2 | 5 (3,4) |
| Anthony Cognato | Vietnam | - | 2 (E07, E12) | 1 | 0.0071 | 1 | 2 (9) |
| Genbank (MN620070.1) | Vietnam | - | 1 (E11) | - | - | 1 | - |
| Anthony Cognato | Vietnam | Cát Tiên Nat. Park | 2 (D09) | 0 | 0 | 1 | 5 (8) |
| Genbank (MN620073.1) | Vietnam | Cát Tiên National Park | 1 (D08) | - | - | 1 | - |
| Anthony Cognato | Vietnam | - | 1 (E15) | - | - | 1 | 4 (9) |
| Genbank (MN620078.1) | Vietnam | - | 1 (E08) | - | - | 1 | - |
| Anthony Cognato | Vietnam | - | 2 (E09) | 0 | 0 | 1 | 3 (9) |
| Genbank (MN620072.1) | Vietnam | - | 1 (E09) | - | - | 1 | - |
| UFFE (14645) | Vietnam | Tam Dao | 15 (E04, E05, E13, E14) | 0.467 | 0.0041 | 1 | 5 (9) |

H2>DNA extraction

To reduce expected contamination by symbiotic and non-symbiotic fungi, specimens first had their mycangia removed, were then washed with 70% alcohol and were cleaned with a paintbrush. DNA was then individually extracted using the Macherey-Nagel NucleoSpin Tissue kit following the manufacturer’s instructions except with two successive elutions in 50 μL BE buffer, and then stored at -18°C.

### Mitochondrial DNA sequencing and statistical analyses

We sequenced 188 specimens, 72 from the native area and 116 from the invaded range (Table 1, Supplementary Table 1), using between one and 16 insects per location. Specimens obtained from Japan originated from the same populations as in Ito and Kajimura (2009). We amplified the barcode COI fragment via PCR using the primers HCO2198 (5’ –TAAACTTCAGGGTGACCAAAAAATCA – 3’) and LCO1490 (5’ – GGTCAACAAATCATAAAGATATTGG – 3’) (Folmer et al. 1994). The PCR was performed as follows: denaturation for 5 min at 94°C followed by 35 cycles of amplification of 45 sec at 94°C, 50 sec at 47°C and 90 sec at 72°C and finally 5 min at 72°C. PCR products were cleaned using the NucleoSpin Gel and PCR Cleanup kit (Machery-Nagel) and sequenced in both directions using the ABI Prism BigDye Terminator v3.1 Cycle Sequencing Kit on an ABI Prism 3500 Genetic Analyzer (Thermo Fisher Scientific). We used CodonCode (CodonCode Corporation) to check electropherograms, create contigs and trim all sequences to 567 bp. DNA sequences were aligned using ClustalW in MEGA X (Kumar et al. 2018). We completed the alignment with all of the barcode COI sequences publicly available from Genbank and for which location information was available, i.e., 32 sequences from 24 localities in 11 countries, including sequences from Cognato et al. (2020) (MN620070.1-MN620078.1), Dole et al. (2010) (GU808708.1-GU808711.1), Ramage et al. (2017) (KX055192.1-KX055198.1), Landi et al. (2017) (KX685266.1), Sire et al. (2019) (MN182983.1, MN183016.1) and Nel et al. (2020) (MT230099.1, MT230101.1, MT230103.1, MT230104.1). The final alignment included 220 individuals.

### Mitochondrial data statistical analysis

We calculated Kimura 2 Parameters (K2P) and p genetic distances between haplotypes and clusters using MEGA X (Kumar et al. 2018). Haplotype and nucleotide diversities were calculated using the pegas package (Paradis 2010) in the R Software (R Core Team 2018). We reconstructed a phylogeny between haplotypes using Maximum-Likelihood and Bayesian inference, with *Xylosandrus germanus* and *X. morigerus* (accession numbers NC_036280.1 and NC_036283.1, respectively). A Maximum-Likelihood phylogeny was performed with MEGA X (Kumar et al. 2018) with 1000 bootstraps using K2P distances. A Bayesian inference of the haplotype phylogeny was performed with MrBayes (Ronquist et al. 2012) with a GTR + I + Γ evolutionary model, and 4 chains run 4 times during 2,000,000 generations with a diagnostic every 100 generations. A median-joining network was realized with PopArt (Bandelt et al. 1999). Lineage and haplotype maps were performed using the R packages maps (Becker et al. 2018), ggplot2 (Wickham 2016) and scatterpie (Guangchuang 2020).

### RAD sequencing and bioinformatics

DNA quantity and quality were assessed using the Qubit dsDNA HS Assay Kit with a Qubit fluorometer. As previously found for the closely related species *Xylosandrus compactus* (Urvois et al. 2022), the DNA amount extracted here from each individual of *Xylosandrus crassiusculus* was too small to allow the direct construction of RAD libraries, therefore we carried out whole genome amplification. We used the Genomiphi kit V3 following the manufacturer’s procedure and the protocol used by Cruaud et al. (2018). Individual RAD libraries were then constructed following Baird et al. (2008) and Etter et al. (2011) with a few modifications listed hereafter. DNA was digested using 250 ng of DNA in 22 μL per sample and 0.5 μL of the PstI-HF enzyme for a total volume of 25 μL. The digested fragments from each specimen were tagged with a unique 5- or 6-bp barcode and a P1 adapter using 1.5 μL of P1 adapter (100 nM) and 0.5 μL of T4 Ligase (2.000.000 U/ml) for a total volume of 30.5 μL. Specimens were then pooled 32 by 32 to create seven libraries. Libraries were sonicated on a Covaris S220 (duty cycle 10%, intensity 5, 200 cycles/burst, duration 75 s) to obtain 300-600 bp fragments. Each library was then tagged with a 5- or 6-nucleotide barcode and a P2 adapter using 1 μL of P2 adapter (10 nM) and 0.5 μL of Quick Ligase (2,000,000 U/ml). The sizing and purification steps were realized using AMPure XP beads (Agencourt). We performed 5 PCR enrichment with 15 cycles (30 ng DNA input, NEB Phusion High-Fidelity PCR Master Mix) for each library to increase fragment diversity. After quality control using the Agilent 2100 Bioanalyzer, the libraries were pooled altogether at an equimolar ratio and sent to MGX-Montpellier GenomiX for sequencing. The library was verified on a Fragment Analyser (Agilent, HS NGS fragment Kit), quantified by qPCR (Kapa Library quantification kit) and sequenced on a SP lane in paired-end 2 × 150 nt mode on a Novaseq6000 (Illumina) according to the manufacturer’s instructions.

We used the RADIS pipeline (Cruaud et al. 2016) to demultiplex individuals using process_radtags (Catchen et al. 2013) and to remove a few low-quality bases at the 3’-ends by trimming all reads at 139 bp. Some specimens were not used in the analysis due to the poor quality of the sequences, leaving between 1 and 7 insects per location for a total of 206 specimens, 81 from the native area and 125 from the invaded range (Table 1, Supplementary Table 1). We obtained on average 7,162,285 (2,534,799 SD) reads per specimen after demultiplexing. We then removed PCR duplicates using clone_filter (Catchen et al. 2013), decreasing the number of reads per individual to 2,323,533 (801,285 SD) on average. To remove potential human DNA contaminations (which can occur during whole genome amplification), we aligned the remaining reads on the reference genome GRCh38.p13 of *Homo sapiens* (accession number: GCA_000001405.28) using the BWA-MEM algorithm (Li and Durbin 2009) and we removed the 103,074 mapping reads, thereby keeping 2,220,459 (785,925 SD) reads per specimen on average.

The following steps were performed using STACKS (Catchen et al. 2013, Rochette et al. 2019) on the Genotoul Bioinformatics Platform (INRAE, Toulouse, France). We ran a pre-analysis on 20 specimens selected to encompass a maximum of genomic diversity to assess the effect of 9 combinations of M and n parameters in STACKS modules: M = 6, 8 and 10 in ustacks (i.e. the maximum distance allowed between stacks) and n = 4, 6 and 8 in cstacks (i.e. the number of mismatches allowed between sample loci when building the catalogue). To remove potential fungal contaminations, we then aligned the obtained loci on the *Ambrosiella xylebori*’s reference genome (accession number: ASM277803v1 (Vanderpool et al. 2018)), as it was the closest complete reference genome available to *X. crassiusculus*’ symbiotic fungus. We used the BWA-MEM algorithm (Li and Durbin 2009) to create a list of potential fungal reads, mapping on the *Ambrosiella xylebori*’s genome, to be later removed in STACKS’ *populations* module. Between 0.14% and 0.16% of the sequences mapped on the *Ambrosiella xylebori*’s genome, depending on the combinations of the parameters M and n, and were all listed to be removed. In STACKS’ *populations* module, we used three possible filtering values for parameter r (the minimum percentage of individuals required to process a locus, here with one population), r = 0, 0.3 and 0.7, respectively. We excluded loci obtained with a mean read depth lower than 8 using VCFtools (Danecek et al. 2011) to test the effect of depth filtering. We compared the number of SNPs and the individual mean depth, homozygosity and missing data for each M, n and r combination. For each combination, we also performed clustering with SNPrelate (Zheng et al. 2012) and used the dendextend R package (Galili 2015) to untangle the obtained dendrograms using the “step1side” method and calculate their entanglement using the entanglement function. The combination of the parameters M and n had limited effects on our results. For a given r, each parameter combination of M and n yielded similar numbers of SNPs (Supplementary Table 2), individual mean depth, homozygosity and missing data (Supplementary Figures S1, S2 & S3), and tree topologies were identical for every parameters combination (Supplementary Figures S4 & S5). Following this pre-analysis, we launched the analysis of the complete dataset, using M = 6, n = 4, and r = 0.8, excluding loci matching *X. crassiusculus*’ symbiotic fungi, and removing loci with a mean depth lower than 8. This M and n parameter set corresponded to the parameters used for *X. compactus* in Urvois et al. (2022). We finally obtained a dataset of 83,055 filtered SNPs.

### RAD SNP statistical analysis

We estimated the specimens’ relative ancestry using Admixture (Alexander et al. 2009), with a putative number of populations, K, ranging from 2 to 20 with a 100-fold cross-validation to assess the best K. We then used the pong 1.4.9 software (Behr et al. 2016) to estimate the major mode (using a greedy approach with 300 runs and a similarity threshold value of 0.95), and plotted and mapped the results using the ggplot2 package (Wickham 2016) in the R Software (R Core Team 2018). A Maximum Likelihood tree was generated using RAxML 8.2.21 (Stamatakis 2014). We used the GTRCAT approximation and allowed the program to automatically halt bootstrapping using the bootstrap converge criterion (Pattengale et al. 2010) through the autoMRE option. The tree was visualized using FigTree V.1.4.4 (https://github.com/rambaut/figtree/releases). Besides, a hierarchical clustering tree was built using SNPRelate (Zheng et al. 2012) on an individual dissimilarity matrix (Zheng 2013). We also calculated, using the StAMPP package (Pembleton et al. 2013), the pairwise Fst (Wright 1951, Weir and Cockerham 1984) and Nei distances (Nei 1972) between the different genomic groups previously obtained with Admixture.

## Results

### Mitochondrial diversity and differentiation

We obtained 50 mitochondrial haplotypes worldwide (Table 1, Figure 1), with 139/567 variable base pairs. Haplotype and nucleotide diversities were 0.954 and 0.074 in the native area and 0.795 and 0.045 in the invaded area, respectively. We found 41 haplotypes in *X. crassiusculus*’ native area, including 16 in Japan, 12 in Vietnam, 11 in China, two in Taiwan, one in India, one in Indonesia and one in Thailand. We found 13 haplotypes in *X. crassiusculus*’ invasive range, including five in Hawaii, four in mainland USA, two in mainland France, American Samoa, French Guiana, French Polynesia, South Africa and Slovenia, and one in Argentina, Canada, Italy, Madagascar, New Zealand, Panama and Spain. Three haplotypes were present in more than four localities (A02, A10 and D02, Figure 1), whereas 36 haplotypes were only found in one locality. Four haplotypes were found in both the native and the invasive areas (A02, A07, A10, D02) (Table 1, Figure 1, Supplementary Table 1). Haplotype A02 was the most widespread haplotype, found in 26 localities in 8 countries (Japan in the native range, and Papua New Guinea, Argentina, Canada, Italy, New Zealand, Spain and the USA in the invasive area). Haplotype D02 was identified in 12 localities in six countries (Indonesia and Japan in the native area, and Papua New Guinea, Madagascar and the Pacific Islands of France and the USA in the invasive area). Haplotype A10 was found in 14 localities in four countries (China, France, Slovenia and the USA), and haplotype A07 was found in two localities in Japan and the USA. The median-joining network showed that five haplotypes (A01, A03, A08, D03 and D04) that were private to the invasive range had a single mutational step from haplotypes found in the native range (Supplementary Figures S6 & S7). Haplotype D10 was three mutational steps away from D09. The haplotype A11 was the closest to the three remaining invasive haplotypes A05, A06, and A09 with 3, 4 and 7 mutational steps, respectively.

**Figure 1:**
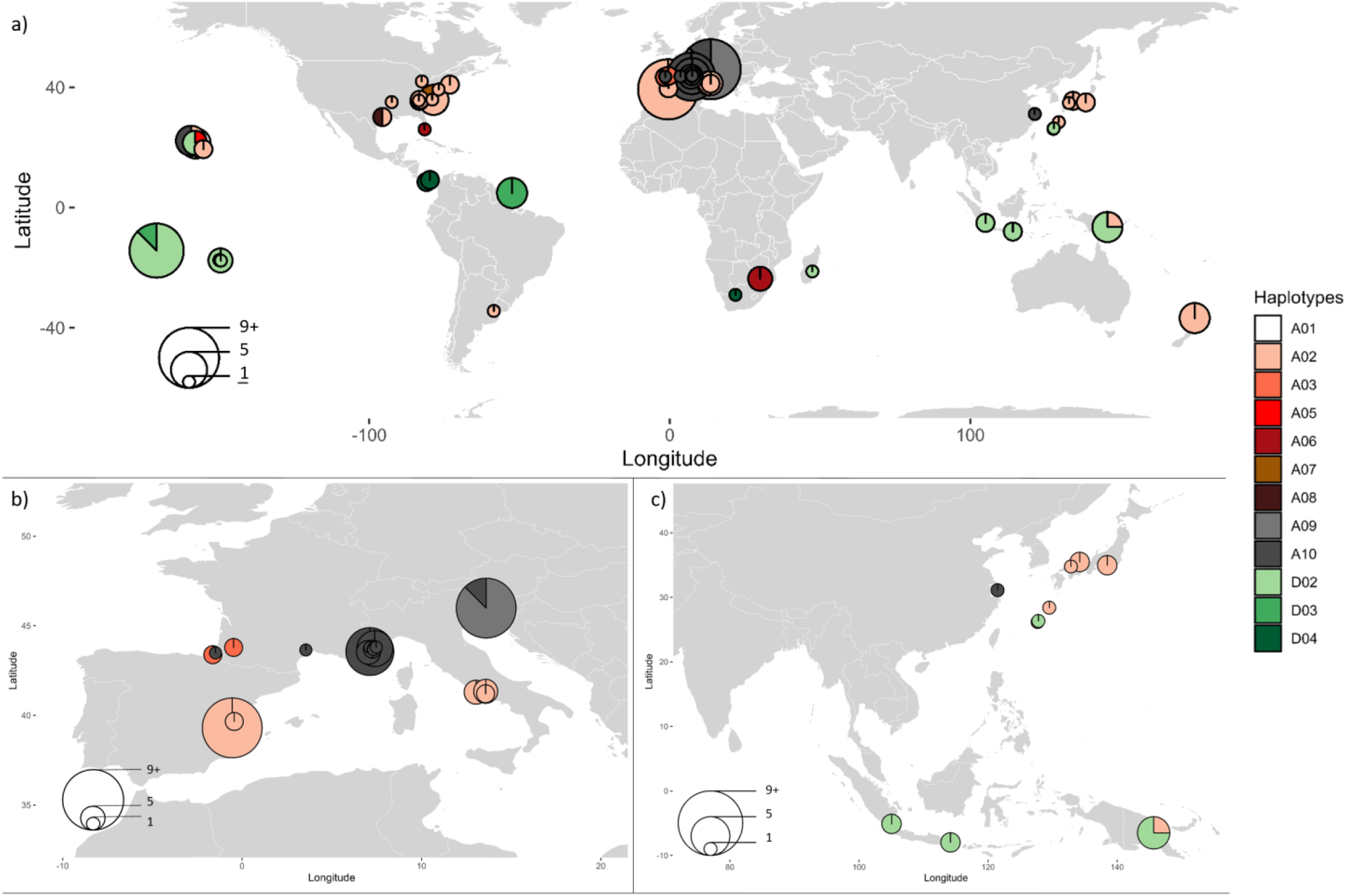
X. crassiusculus’ *invasive haplotype map a) worldwide; b) focusing on Europe, and c) and focusing on Asia. The coordinates of some localities were not known (cf. Table 1). Their pies were thus added at approximate coordinates.*

The Maximum-Likelihood and the Bayesian inference trees suggested that haplotypes can be grouped in two differentiated Clusters (Figure 2, Supplementary Figures S8 & S9) generally consistent with the results found by Storer et al. (2017). The average K2P and p genetic distances within Clusters were 0.054 (0.032 Standard Error Estimate) and 0.050 (0.010 SEE), and between Clusters of 0.118 (0.008 SEE) and 0.109 (0.010 SEE), respectively (Supplementary Table 3). Both Clusters were further structured into five mitochondrial lineages, Cluster 1 including lineages D and E, and Cluster 2 including lineages A, B and C (Table 1, Figures 3, Supplementary Figure S8). To ease the comparison of results across studies, our lineages A, B and C fully correspond to the lineages A, B and C already identified in Ito and Kajimura’s study (2009), and we named D and E the two new mitochondrial lineages we identified in the present work. The K2P distances within lineages were lower than 0.049 (mean 0.020) (Table 1) and between lineages higher than 0.070 (mean 0.103) (Table 2). Lineage A comprised 12 haplotypes and was present in the native area in China and Japan, and in the invaded area in France, Italy, Slovenia, Spain, Argentina, New Zealand, Papua New Guinea, Canada, mainland USA and Hawaii (Figure 3). Lineages B and C were exclusively found in Japan and consisted of 7 and 6 haplotypes, respectively. Lineage D comprised 10 haplotypes and was found in the native area in China, India, Indonesia, Japan, Thailand, Vietnam, and in the invaded area in Papua New Guinea, Madagascar, Panama, French Polynesia, Hawaii, American Samoa and French Guiana. Finally, lineage E was composed of 15 haplotypes and was found exclusively in *X. crassiusculus*’ native area in China, Taiwan and Vietnam. The Maximum-Likelihood tree reached high support (>0.50) for most nodes except for some nodes within lineage A and within lineage E (Figure 2). On the other hand, the Bayesian inference tree had a lower resolution and had low support for the separation between lineages A and B (Posterior Probability = 0.78) but high support for lineages C, D, and E (Supplementary Figure S9).

**Table 2:** Genetic distances within and between COI lineages (derived from the mitochondrial haplotypes) and their respective standard error estimates based on the Kimura 2-parameter model. The genetic distances between COI lineages are on the bottom left part, and the standard error estimates on the top left part of the table.

|  | Within |  | Between |  |  |  |  |
| --- | --- | --- | --- | --- | --- | --- | --- |
|  | Distances | Standard Error Estimates | Lineage A | Lineage B | Lineage C | Lineage D | Lineage E |
| Lineage A | 0.0135 | 0.003 |  | 0.011 | 0.0117 | 0.0123 | 0.014 |
| Lineage B | 0.0076 | 0.0024 | 0.0709 |  | 0.0128 | 0.0135 | 0.0147 |
| Lineage C | 0.014 | 0.0039 | 0.0798 | 0.0932 |  | 0.0135 | 0.0145 |
| Lineage D | 0.0482 | 0.0063 | 0.1095 | 0.1242 | 0.1158 |  | 0.01 |
| Lineage E | 0.0152 | 0.0029 | 0.1176 | 0.1252 | 0.1188 | 0.0783 |  |

**Figure 2:**
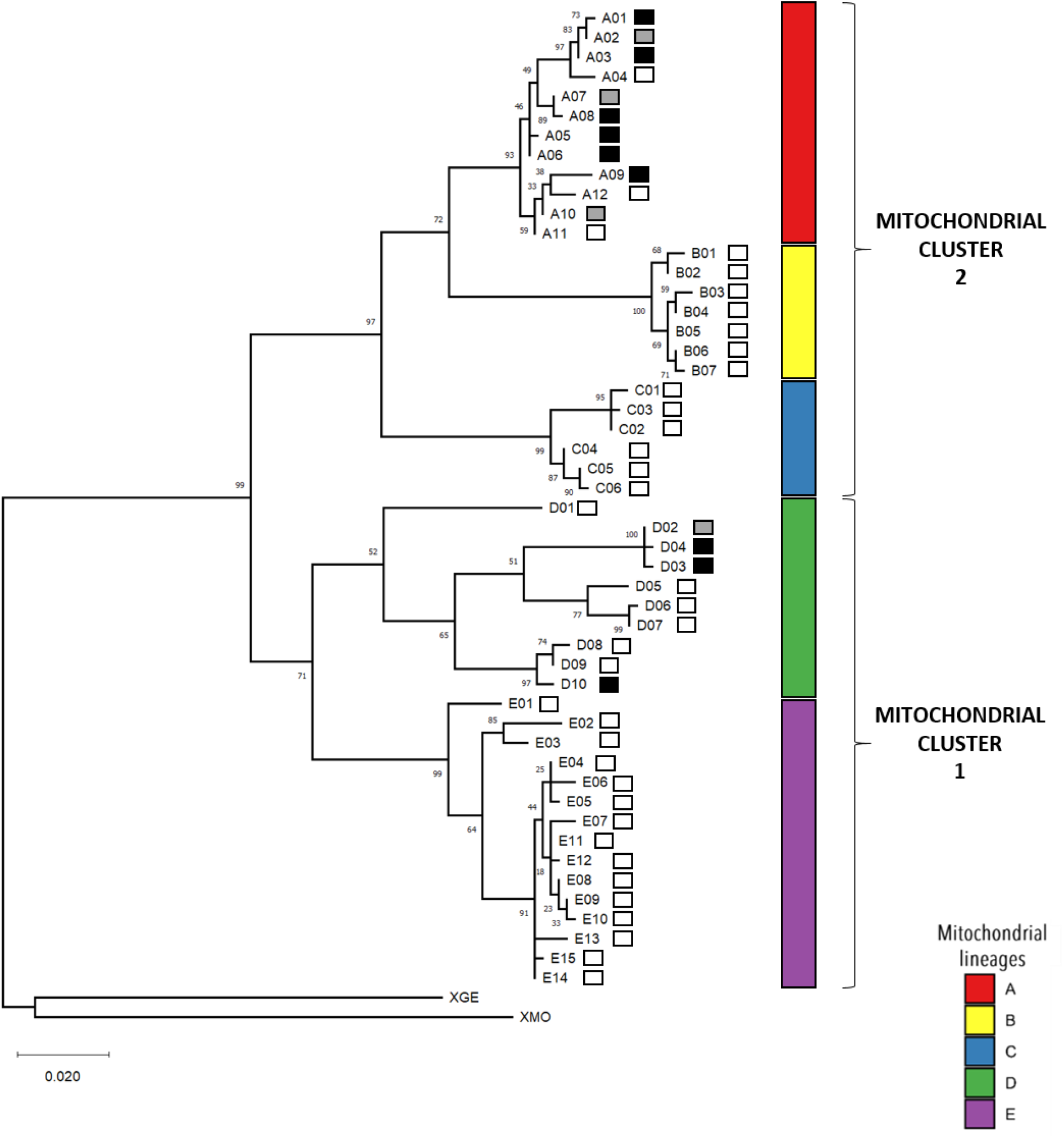
*Maximum Likelihood tree based on* X. crassiusculus’ *COI sequences built with MEGA X. We used* Xylosandrus germanus *(XGE NC036280.1) and* Xylosandrus morigerus *(XMO NC_036283.1) as outgroups. The color of the squares next to the haplotypes shows whether they were identified in the native area only (white), in the invaded area only (black) or in both areas (grey).*

**Figure 3:**
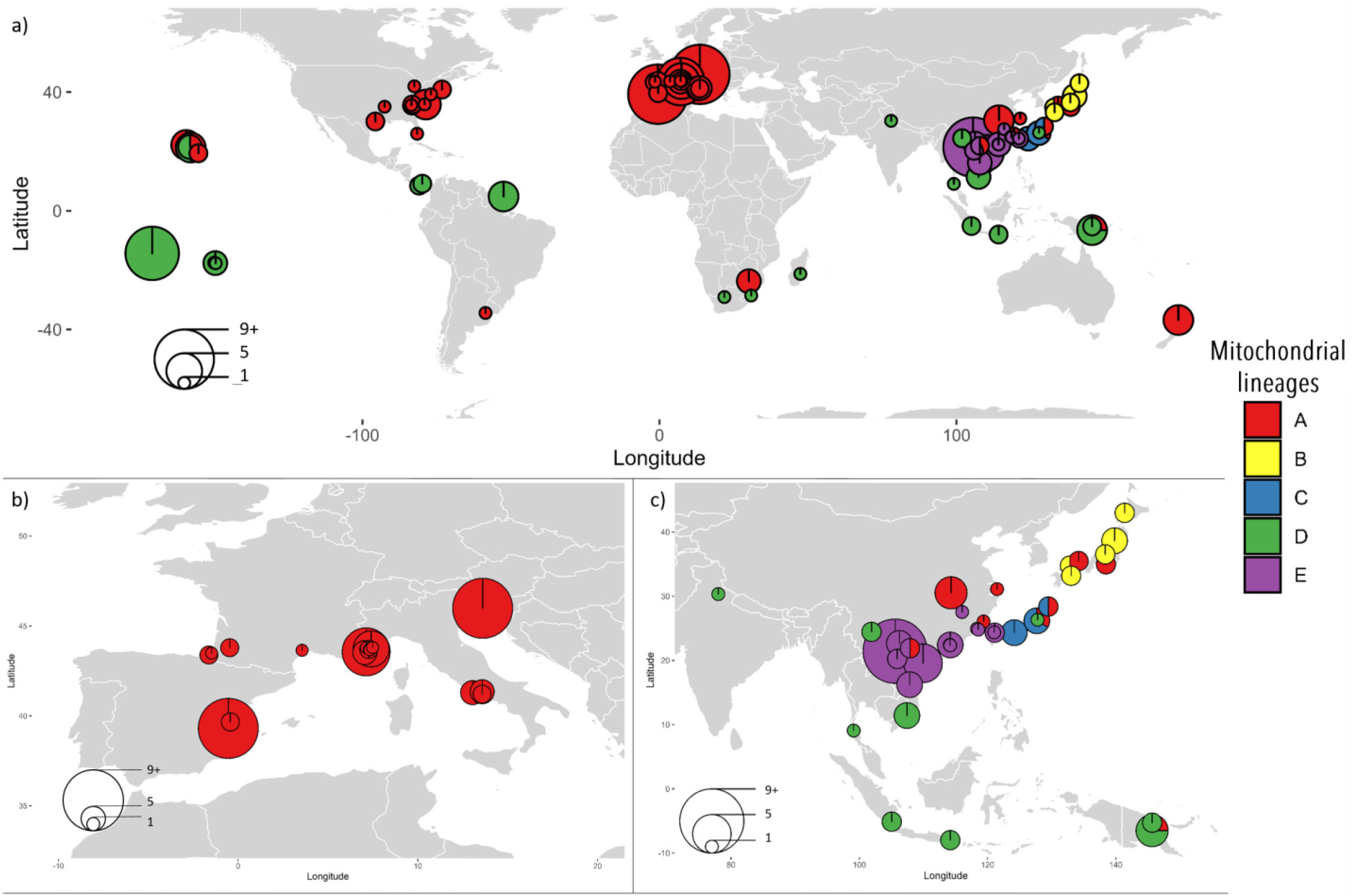
X. crassiusculus’ *mitochondrial lineage map a) worldwide ; b) focusing on Europe, and c) focusing on Asia. The coordinates of some localities were not known (cf. Table 1). Their pies were thus added at approximate coordinates. The mitochondrial lineages were derived from mitochondrial the haplotypes.*

### Genetic structure at nuclear SNPs obtained from RAD sequencing

The average homozygosity estimated from RADseq data was 0.993 (0.016 SD), and the average inbreeding coefficient was 0.958 (0.101 SD). When running Admixture on the complete dataset, the cross-validation values reached a plateau for K = 9, which we considered the most parsimonious number of genomic groups (Supplementary Figure S10). With a similarity threshold of 0.95, the 300 Admixture runs yielded 82 modes, the major mode representing 109 runs with a pairwise similarity of 0.978.

The genomic groups obtained at K = 2 matched the mitochondrial Clusters 1 and 2 for most individuals, except for 17 specimens showing a low assignation score and corresponding to the mitochondrial lineages B and C (Supplementary Figure S11) from Japan. These particular specimens formed the third genomic group at K = 3 (Figure 4), the two other genomic groups corresponding to (i) all the individuals of the mitochondrial Cluster 1, and (ii) the individuals of Cluster 2 restricted to the lineage A described above (Figure 4). We will hereafter refer to these three groups as Cluster JapB-C and genomic Clusters 1 and 2 (Figure 4). For the optimal K = 9, the Cluster JapB-C was further split into the genomic groups 1 and 2 (tightly corresponding to the mitochondrial lineages B and C mentioned above), genomic Cluster 2 was composed of the genomic groups 3, 4 and 5 (including all individuals from the mitochondrial lineage A) and genomic Cluster 1 comprised the genomic groups 6, 7, 8 (corresponding to mitochondrial lineage D) and 9 (mitochondrial lineage E).

**Figure 4:**
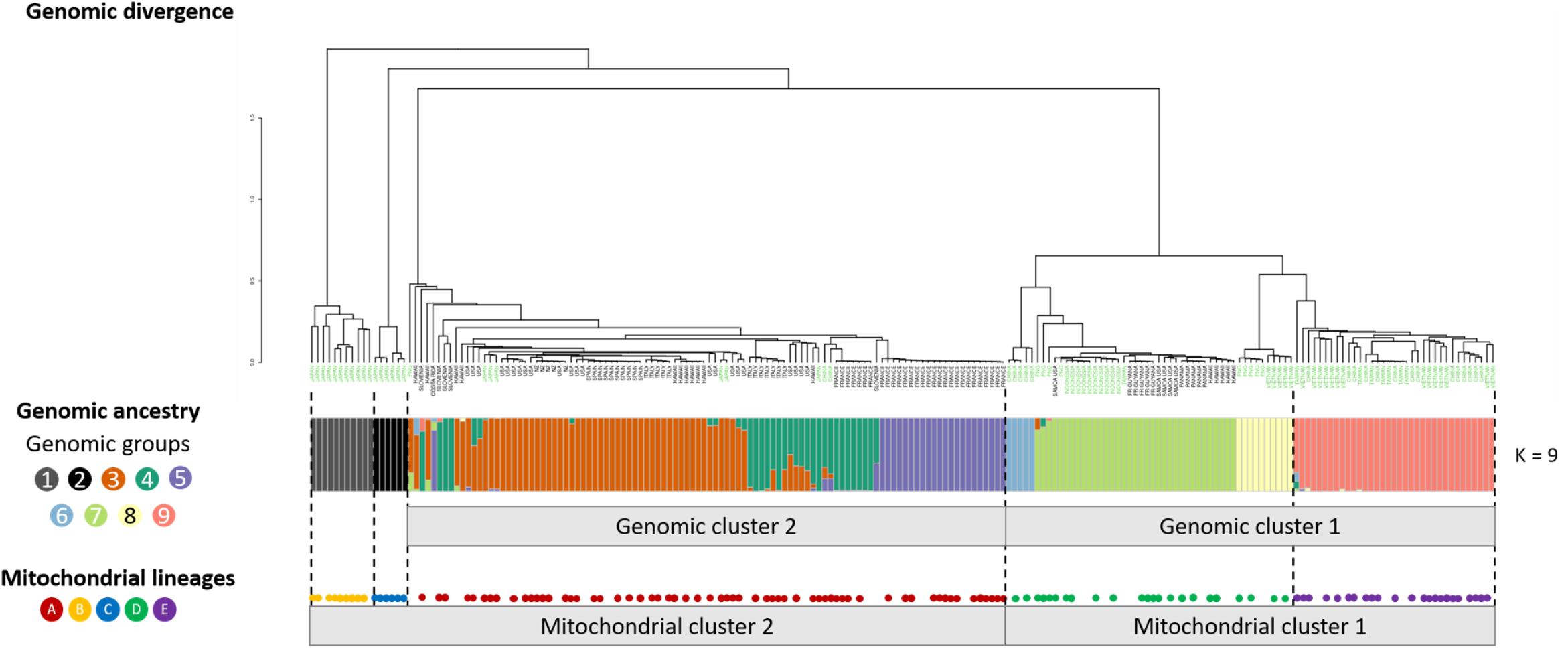
*Clustering tree and admixture plot for K = 9 calculated on* Xylosandrus crassiusculus’ *RAD sequencing data. The leaves labels of the clustering tree represent the country of origin of the samples, and the countries in the* X. crassiusculus’ *native area are represented in green. When available, the specimens’ mitochondrial lineage, derived from the mitochondrial haplotypes, are represented as colored circles at the bottom of the figure.*

For K = 9, 177 specimens were assigned to their genomic group with a score higher than 0.95, and 154 scored more than 0.999. Within genomic Cluster 1, two groups were restricted to the native area (genomic group 6 occurred only in one locality in China and genomic group 9 was present in Taiwan, Vietnam and China (Figure 5)), and two groups were invasive in different regions of the world (genomic group 7 was native from Indonesia and invasive in the Pacific Islands, Costa Rica, French Guiana and Papua New Guinea; genomic group 8 was native from Vietnam and invasive in Papua New Guinea and Vietnam). Within genomic Cluster 2, the three identified genomic groups were found both in native and invasive areas. Group 3 was found in Japan (native area) and Hawaii, mainland USA, Spain, Italy, New Zealand and Papua New Guinea in the invaded area (Figure 6). Genomic group 4 was found in Japan and China in the native range, and in Italy, France, Slovenia,, Hawaii and mainland USA in the invasive regions. Note that some specimens from the USA were assigned as admixed between genomic groups 3 and 4. Genomic group 5 was composed exclusively of specimens from France and one specimen from Costa Rica, and was thereby not found in the native range.

**Figure 5:**
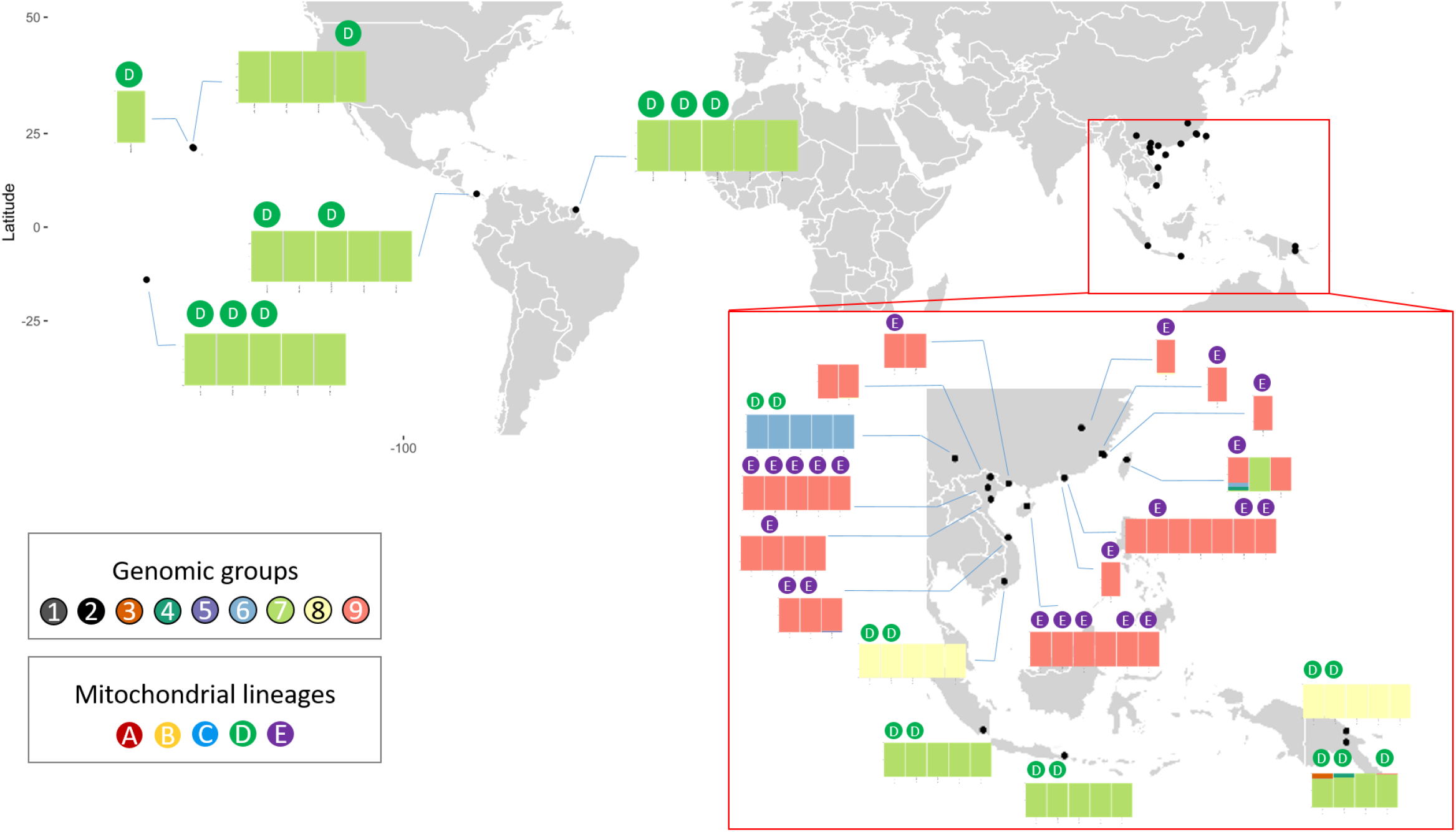
Map representing the admixtures plot for the specimens belonging to the genomic cluster 1. The specimens’ mitochondrial lineage, derived from its mitochondrial haplotype, was added on top of the barplots when available.

**Figure 6:**
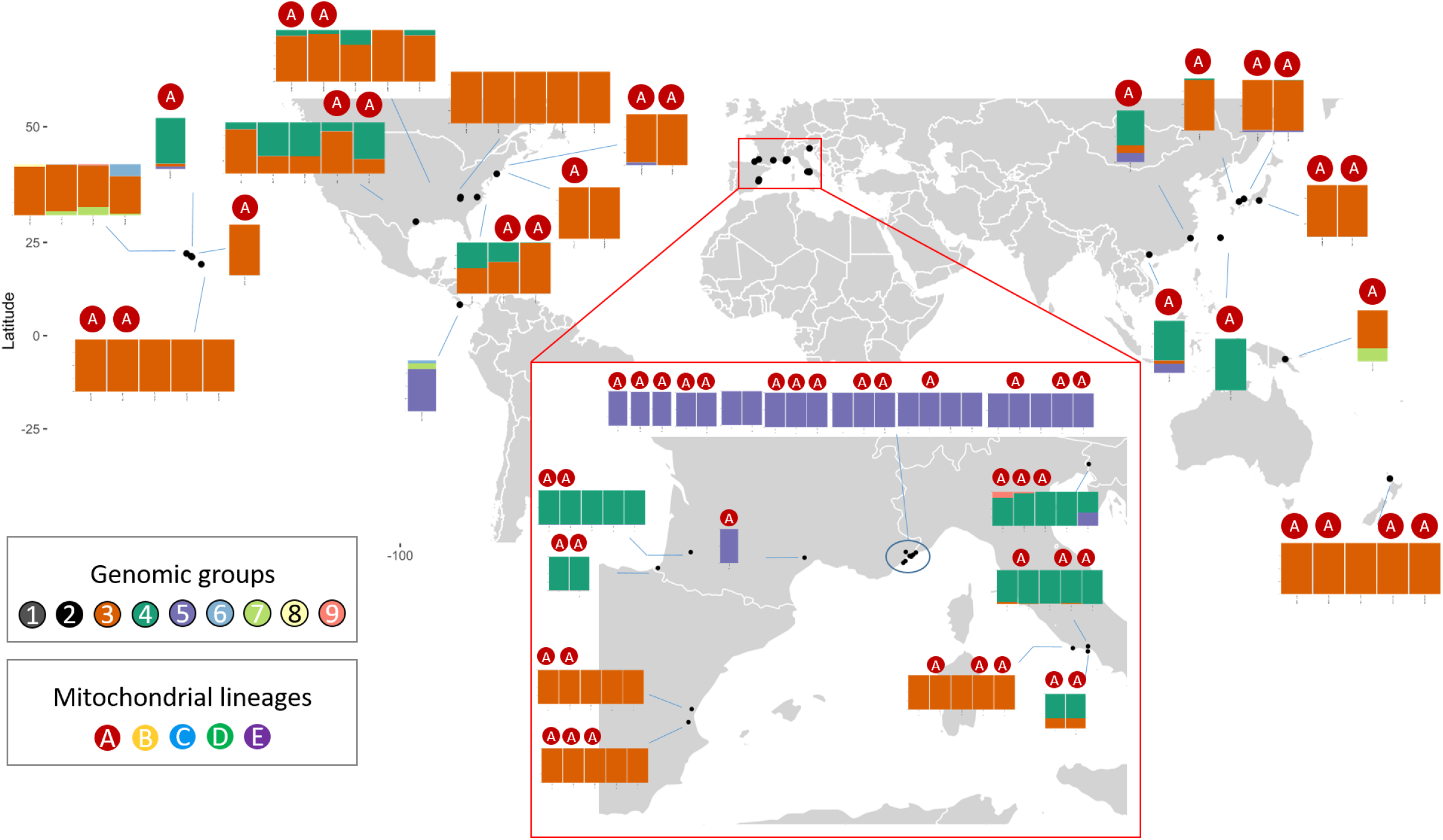
Map representing the admixtures plot for the specimens belonging to the genomic cluster 2. The specimens’ mitochondrial lineage, derived from its mitochondrial haplotype, was added on top of the barplots when available.

The Fst and Nei distances between genomic Clusters were 0.956 and 0.366, respectively (Table 3). Between genomic groups, they averaged 0.894 (0.125 SD) and 0.294 (0.132 SD), respectively. Our results showed a lower divergence between genomic groups within Cluster 2 than within Cluster 1, with average Fst of 0.529 (0.0132 SD) and 0.847 (0.059 SD), and Nei distances of 0.0189 (0.005 SD) and 0.102 (0.020 SD), respectively.

**Table 3:** Fst (bottom left part) and Nei (top left part) distances between genomic groups.

|  | Genomic<br>group 1 | Genomic<br>group 2 | Genomic<br>group 3 | Genomic<br>group 4 | Genomic<br>group 5 | Genomic<br>group 6 | Genomic<br>group 7 | Genomic<br>group 8 | Genomic<br>group 9 |
| --- | --- | --- | --- | --- | --- | --- | --- | --- | --- |
| Genomic group 1 | 0 | 0,403 | 0,383 | 0,375 | 0,38 | 0,368 | 0,373 | 0,384 | 0,37 |
| Genomic group 2 | 0,899 | 0 | 0,328 | 0,321 | 0,327 | 0,381 | 0,382 | 0,388 | 0,381 |
| Genomic group 3 | 0,948 | 0,96 | 0 | 0,013 | 0,024 | 0,358 | 0,364 | 0,37 | 0,363 |
| Genomic group 4 | 0,907 | 0,923 | 0,411 | 0 | 0,019 | 0,362 | 0,375 | 0,371 | 0,364 |
| Genomic group 5 | 0,939 | 0,969 | 0,671 | 0,504 | 0 | 0,359 | 0,369 | 0,371 | 0,363 |
| Genomic group 6 | 0,913 | 0,957 | 0,965 | 0,94 | 0,977 | 0 | 0,075 | 0,126 | 0,108 |
| Genomic group 7 | 0,943 | 0,962 | 0,961 | 0,949 | 0,967 | 0,855 | 0 | 0,12 | 0,102 |
| Genomic group 8 | 0,928 | 0,965 | 0,968 | 0,947 | 0,979 | 0,919 | 0,906 | 0 | 0,081 |
| Genomic group 9 | 0,912 | 0,93 | 0,949 | 0,926 | 0,947 | 0,8 | 0,83 | 0,768 | 0 |

The RAxML analysis stopped after 400 bootstraps with a best tree scoring a likelihood score of - 359,651.47. The phylogenetic tree was very consistent with the clustering tree and Admixture results described above. It revealed two groups corresponding to the Cluster JapB-C (Admixture groups 1 and 2) and two distinct Clusters corresponding to the genomic Clusters 1 and 2, respectively split in 4 and 3 major branches (Supplementary Figure S12) matching groups 6 to 9 on the one hand, and groups 3 to 5 on the other hand.

## Discussion

The aim of this study was to analyze the genetic structure and the invasion history of *X. crassiusculus* worldwide, using a strategy that reconciled previous studies (which used different markers and focused on different regions) and to obtain a global picture including previously unstudied areas such as Europe. To do so, we sequenced the barcode mitochondrial marker in all regions, which allowed us to merge the data previously published from South America and South Africa and sequences retrieved from GenBank. We included a subset of the individuals studied in Ito & Kajimura (2009) corresponding to all the clades and subclades they identified using the second half of the COI gene (i.e., a fragment which did not overlap with the barcode region), which allowed for an interpretation of our results in the light of the Japanese genetic structure. Finally, we also included specimens from the main regions studied in Storer et al. (2017) which allowed for an evaluation of the consistency between their results and ours. This sampling design and the use of both mitochondrial and nuclear pangenomic markers provided a broader view of the genetic structure and of the dispersal around the globe.

To facilitate the comparisons between studies, we named the main genetic groups and lineages consistently with previous results, in particular our mitochondrial clusters A, B and C corresponded to the clusters identified by Ito and Kajimura (2009) under the same codes. Both mitochondrial and nuclear markers highlighted the existence of two main genetic Clusters that were generally consistent with the results found by Storer et al. (2017). The same genetic groups in our study and theirs are consistently named Cluster 1 and Cluster 2.

### Genetic structure in *Xylosandrus crassiusculus*

#### *The consolidated global genetic structure of* X. crassiusculus

Most individuals fall into two main highly differentiated genetic clusters supported both by mitochondrial and nuclear data, in this as well as in previous studies. Interestingly, Clusters are mostly allopatric and overlap only in a few regions (Figure 7). Cluster 1 is native to China, India, Indonesia, Taiwan, Thailand (Storer et al. 2017) and Vietnam, and was found once in Okinawa (Storer et al. 2017, Cognato et al. 2020). It is invasive in Papua New Guinea, Central America, the Pacific islands, and several African countries (Storer et al. 2017, Nel et al. 2020). Cluster 2 is native to China, Japan and Taiwan (Ito and Kajimura 2009, Storer et al. 2017), and invasive in South, Central, and North America (Storer et al. 2017), in the Pacific Islands, Africa (Nel et al. 2020), Oceania and Europe. The two Clusters were only found together in the native area in Taiwan (Storer et al. 2017), the Guangxi province in China and Okinawa. In the invaded range, they co-occurred in O’ahu Island in Hawaii, in Papua New Guinea and in South Africa. Both Clusters were also found in Central America, but each in a different country. Moreover, their worldwide distributions suggest that Cluster 1 had a circumtropical distribution while Cluster 2 was present at higher latitudes and in temperate regions. The apparent difference between the two Clusters’ geographical distributions may associate with various factors such as different abiotic requirements (i.e., climatic requirements), different biotic interactions (i.e., host association) or different colonizable ranges due, for instance, to different means of passive transportation (Guisan et al. 2017). Tests of these hypotheses will require additional sampling and specific analyses.

**Figure 7:**
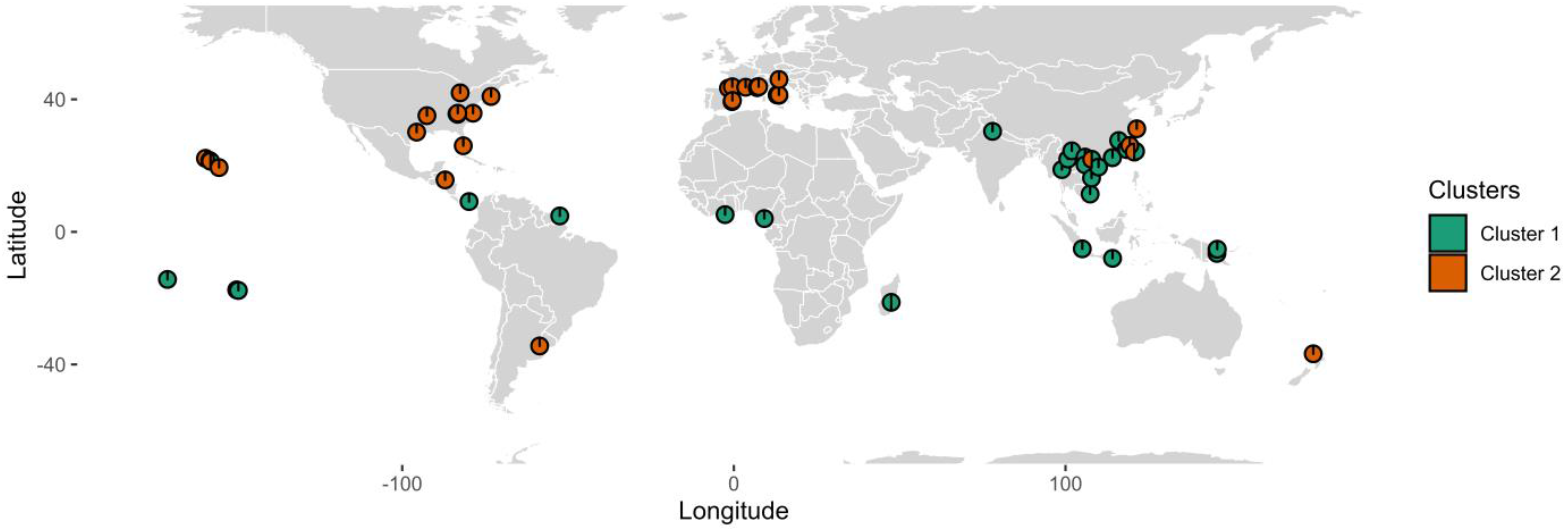
*Map representing the worldwide distribution of the two genomic Clusters of* X. crassiusculus.

#### Mitonuclear discordance in one clade found in Japan

Most specimens could be unambiguously assigned to one of the two main genetic Clusters discussed above. Still, a group of 17 individuals found exclusively in Japan (mitochondrial lineages B and C, this study and Ito & Kajimura (2009)) were placed in a third genomic Cluster we called JapB-C. However, these individuals were phylogenetically close to the ones belonging to mitochondrial lineage A and were thus expected to belong to Cluster 2. Such a discrepancy between markers is referred to as a case of mitonuclear discordance. It can result from diverse phenomena, including natural selection, introgression, sex-bias in offsprings, sex-biased dispersal (El Mokhefi et al. 2016) or reproductive manipulation by *Wolbachia* (Sloan et al. 2017). Still, it is beyond the scope of this study to investigate the cause of the observed differences.

#### *Xylosandrus crassiusculus*’ invasion history and pathways

Our study is the first to include specimens from Europe. Europe was only invaded by Cluster 2, with specimens from the three genomic groups (3, 4 and 5), and four haplotypes from the mitochondrial lineage A. The closely related species *X. germanus* (Dzurenko et al. 2020) and *X. compactus* (Urvois et al. 2022) seem to have spread across Europe from single introductions, but *Xylosandrus crassiusculus*’ likely invaded Europe multiple times from multiple sources, followed by local dispersal as observed for other xyleborine species in North America (Cognato et al. 2015, Smith and Cognato 2022). The first invaded Italian locality among our sampling sites (Circeo National Park) is characterized by haplotype A02 and the genomic group 3. This could point to a source in Japan if the species was directly introduced from the native range, or to a source in the US, as several North American localities invaded during the 20th century had the same genetic characteristics (Figure 8). Spanish populations were genetically very similar to the Italian ones, suggesting that *X. crassiusculus* colonized Spain directly from the Italian source through intracontinental movements. Bridgehead events from the USA to Europe were already documented for various invasive insect species, for example, *Harmonia axyridis* (Lombaert et al. 2010), *Leptoglossus occidentalis* (Lesieur et al. 2018) or *Anoplophora glabripennis* (Javal et al. 2019).

**Figure 8:**
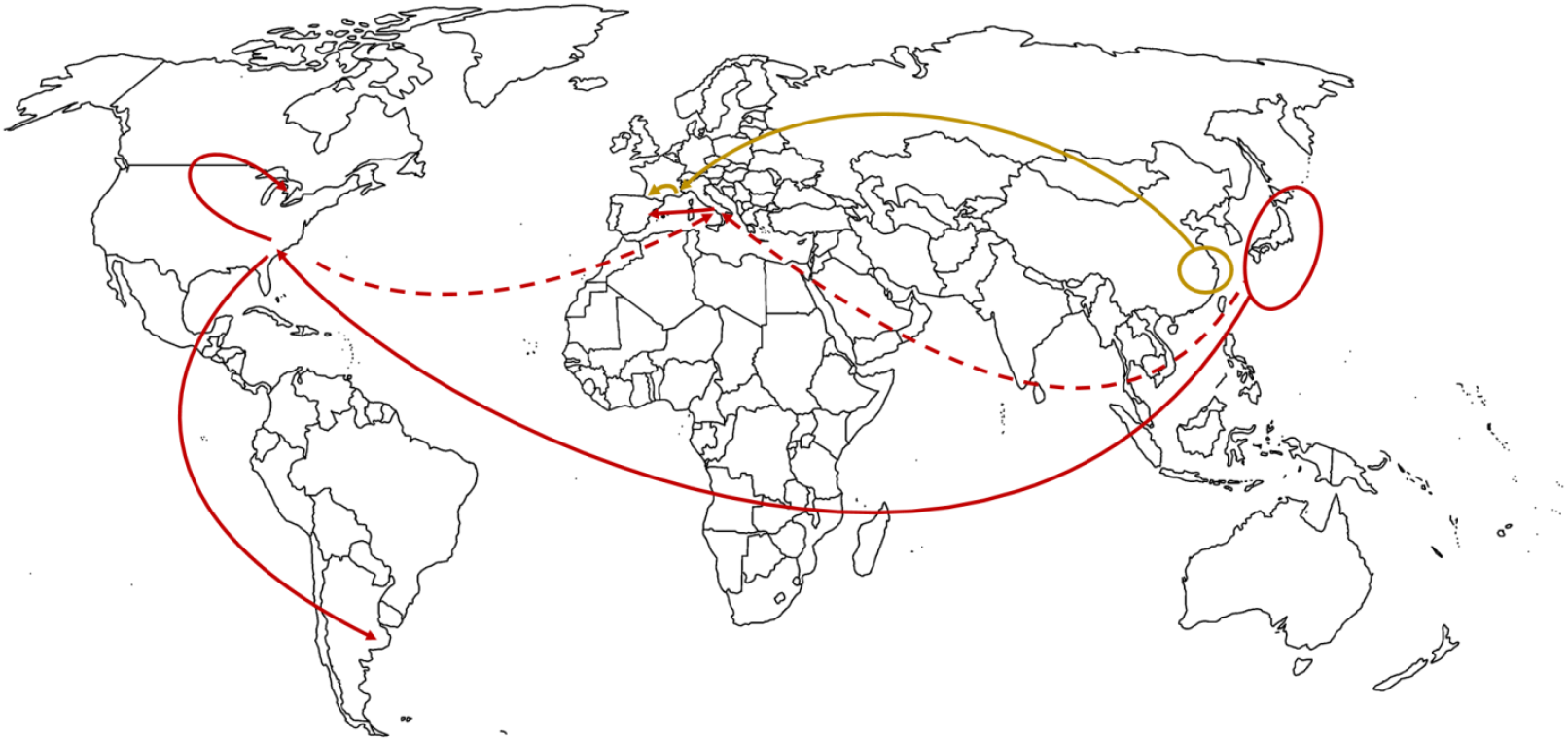
Potential invasion scenarios of Xylosandrus crassiusculus in the USA, Canada, Argentina and Europe.

Specimens from Southeastern France bore haplotype A10, also found in Shanghai in the native area. This suggests an independent colonization event from the region of Shanghai, similar to *X. compactus* (Urvois et al. 2022). However, as Shanghai is one of China’s most economically important cities and the busiest port worldwide (UNCTAD 2020), it could also have acted as a bridgehead by exporting infested plants coming from other areas in China or other countries in Asia, as observed for the invasive box tree moth *Cydalima perspectalis* by Bras et al. (2019). A few individuals with similar genetic characteristics were found in various European localities where *X. crassiusculus* was later detected, as in Eastern Slovenia, Southern France and Southwestern France, suggesting stepping-stone expansion from Southeastern France to nearby regions. Other European specimens could correspond to other independent colonizations, such as specimens belonging to genomic group 4 with haplotype A02 in Italy or A03 in Southwestern France, or specimens with haplotype A9 in Slovenia. These genetic combinations were not identified in the native or invaded areas, we thus cannot infer the source of these populations.

Our results also brought complementary information to document the invasion history of *X. crassiusculus* in the Americas and Africa. Despite extensive sampling, Storer et al. (2017) exclusively found Cluster 2 in the USA, which agrees with our results. The presence of genomic group 3 and haplotypes A02 and A07 in the USA suggests that Japan could be the donor area, similar to the introduction of *X. germanus* to the USA (Dzurenko et al. 2020). Several localities in the USA across different states were genetically similar, and the same mitochondrial haplotype (A02) was found in localities in the USA, Canada and Argentina. We thus hypothesize that mainland USA could have acted as a source for further expansion to Canada and Argentina through stepping-stone expansion and bridgehead events. Storer et al. (2017) reported Africa was invaded from mainland Asia (Cluster 1) and suggested that historic dispersal could explain the high differentiation between Madagascar and other African locations. Our analysis showed that the haplotypes reported by Nel et al. (2020) in South Africa belonged to both Clusters. A similar situation was reported for *Euwallacea fornicatus* with two divergent haplogroups, potentially corresponding to cryptic species, co-occurring in South Africa (Bierman et al. 2022). The seemingly special place of South Africa on the African continent could result from larger imports or traffic due to some of the largest African ports, such as Port Durban or Port of Richards Bay. However, it is also possible that both Clusters also co-occur in other African countries as the sampling in Africa remains very limited. Indeed, *X. crassiusculus* was reported in 15 African countries (Nel et al. 2020), but few specimens were sampled in only four of them.

### Monitoring *Xylosandrus crassiusculus*’ Clusters in the invaded area

Our study confirmed that *X. crassiusculus* comprises at least three differentiated genomic clusters, two of which are invasive worldwide. The strong intraspecific differentiation documented from nuclear and mitochondrial markers and the existence of the Cluster JapB-C suggest a reassessment of potential species limits within *X. crassiusculus*. The p distance between Clusters 1 and 2 based on COI obtained in this study was 10.9% (Standard Error Estimate = 1.0%), fitting in the range reported by Cognato et al. (2020) for the probable recognition of new species of Xyleborini (>10-12% COI and/or >2-3% using the nuclear CAD gene). Other criteria could help rule on the Clusters’ status, such as existing gene flow or differences in their ecology. While most specimens were unambiguously assigned to one genomic Cluster for K = 3, the few specimens partly assigned to both Clusters could suggest existing hybridization. Our study also showed that most regions of the world had only one Cluster, suggesting differences in the Clusters’ ecological preferences.

We call for future research comparing the two Clusters’ biology and ecology. In case of ecological differences, detecting the arrival of a so far non-occurring Cluster in already invaded areas would be crucial information. The newly introduced Cluster could have a different invasion dynamic, establishing in localities with unsuitable conditions for the other Cluster, attacking new host tree species, or having different phenology with a higher voltinism. In turn, surveillance should be maintained in already invaded countries and genetic expertise should be deployed to identify the intercepted specimens at the Cluster scale and possibly detect the arrival of a so far non-occurring lineage. This early detection would also facilitate the implementation of measures to eradicate populations at their earliest stages (LaBonte 2010), or to control and mitigate the damage caused by *X. crassiusculus* if eradication is not possible.

## Acknowledgement

We thank Richard Bellanger, Jared Bernard, Ben Boyd, Massimo Faccoli, Steve Frank, Diego Gallego, Andreja Kavčič and Jhunior Morillo for sending specimens. We thank GenSeq technical facilities of ISEM, Institut des Sciences de l’Evolution de Montpellier, supported by the Labex CeMEB and ANR “Investissements d’Avenir” program (ANR-10-LABX-04-01), for the use of the ultrasonicator. We are grateful to MGX-Montpellier GenomiX for sequencing the RAD libraries and to the Genotoul bioinformatics platform Toulouse Occitanie (Bioinfo Genotoul, doi: 10.15454/1.5572369328961167E12) for providing computing and storage resources. We are also grateful to the Sino-French Joint Laboratory for Invasive Forest Pests in Eurasia (IFOPE) for its support.

## Data accessibility

Individual RAD sequences files are available in a fq.gz format a the Sequence Read Archive (SRA) (Study Accession no. XXXXX). The VCF files XXXXX.vcf as well as the popmap used in STACKS’population modile and specimens metadata (e.g. GPS coordinates) are available on Portail Data INRAE (XXXXX). The Genbank accession numbers for the mitochondrial haplotypes reported in this paper are XXXXX to XXXXX.

## Author contribution statement

Marie-Anne Auger-Rozenberg and Carole Kerdelhué designed the study. Laure Sauné, Claudine Courtin and Teddy Urvois completed the molecular biology work. Teddy Urvois and Charles Perrier performed the bioinformatics, the statistical analysis and made the figures. Alain Roques, Jiri Hulcr, Hisashi Kajimura and Anthony I. Cognato performed the field work and helped interpret the results. Teddy Urvois, Carole Kerdelhué, Marie-Anne Auger-Rozenberg and Charles Perrier wrote the original draft of the manuscript. All authors reviewed, edited and approved the final version of the manuscript.

## Conflict of interst

The authors declare no conflict of interests. Specimens sampled did not involve endangered nor protected species.

## References

Alexander, D. H., J. Novembre, and K. Lange. 2009. Fast model-based estimation of ancestry in unrelated individuals. Genome Research 19:1655–1664.

Anderson, D. M. 1974. First record of Xyleborus semiopacus in the continental United States (Coleoptera, Scolytidae). Cooperative Economic Insect Report 24:863–864.

Baird, N. A., P. D. Etter, T. S. Atwood, M. C. Currey, A. L. Shiver, Z. A. Lewis, E. U. Selker, W. A. Cresko, and E. A. Johnson. 2008. Rapid SNP discovery and genetic mapping using sequenced RAD markers. PLoS ONE 3:e3376.

Bandelt, H.-J., P. Forster, and A. Röhl. 1999. Median-joining networks for inferring intraspecific phylogenies. Molecular Biology and Evolution 16:37–48.

Becker, R. A., A. R. Wilks, R. Brownrigg, T. P. Minka, and A. Deckmyn. 2018. maps: draw geographical maps. R package version 3.3.0.

Behr, A. A., K. Z. Liu, G. Liu-Fang, P. Nakka, and S. Ramachandran. 2016. pong: fast analysis and visualization of latent clusters in population genetic data. Bioinformatics 32:2817–2823.

Bertelsmeier, C., S. Ollier, and X. Liu. 2021. Bridgehead effects distort global flows of alien species. Diversity and Distributions 27:2180–2189.

Bierman, A., F. Roets, and J. S. Terblanche. 2022. Population structure of the invasive ambrosia beetle, Euwallacea fornicatus, indicates multiple introductions into South Africa. Biological Invasions.

Bras, A., D. N. Avtzis, M. Kenis, H. Li, G. Vétek, A. Bernard, C. Courtin, J. Rousselet, A. Roques, and M.-A. Auger-Rozenberg. 2019. A complex invasion story underlies the fast spread of the invasive box tree moth (Cydalima perspectalis) across Europe. Journal of Pest Science 92:1187–1202.

Catchen, J., P. A. Hohenlohe, S. Bassham, A. Amores, and W. A. Cresko. 2013. Stacks: an analysis tool set for population genomics. Molecular Ecology 22:3124–3140.

Cognato, A. I., E. R. Hoebeke, H. Kajimura, and S. M. Smith. 2015. History of the Exotic Ambrosia Beetles Euwallacea interjectus and Euwallacea validus (Coleoptera: Curculionidae: Xyleborini) in the United States. Journal of Economic Entomology 108:1129–1135.

Cognato, A. I., G. Sari, S. M. Smith, R. A. Beaver, Y. Li, J. Hulcr, B. H. Jordal, H. Kajimura, C.-S. Lin, T. H. Pham, S. Singh, and W. Sittichaya. 2020. The essential role of taxonomic expertise in the creation of DNA databases for the identification and delimitation of Southeast Asian ambrosia beetle species (Curculionidae: Scolytinae: Xyleborini). Frontiers in Ecology and Evolution 8.

Cruaud, A., M. Gautier, J. P. Rossi, J. Y. Rasplus, and J. Gouzy. 2016. RADIS: analysis of RAD-seq data for interspecific phylogeny. Bioinformatics 32:3027–3028.

Cruaud, A., G. Groussier, G. Genson, L. Saune, A. Polaszek, and J. Y. Rasplus. 2018. Pushing the limits of whole genome amplification: successful sequencing of RADseq library from a single microhymenopteran (Chalcidoidea, Trichogramma). PeerJ 6:e5640.

Danecek, P., A. Auton, G. Abecasis, C. A. Albers, E. Banks, M. A. DePristo, R. E. Handsaker, G. Lunter, G. T. Marth, S. T. Sherry, G. McVean, R. Durbin, and G. Genomes Project Analysis. 2011. The variant call format and VCFtools. Bioinformatics 27:2156–2158.

Dole, S. A., B. H. Jordal, and A. I. Cognato. 2010. Polyphyly of Xylosandrus Reitter inferred from nuclear and mitochondrial genes (Coleoptera: Curculionidae: Scolytinae). Molecular Phylogenetics and Evolution 54:773–782.

Dzurenko, M., C. M. Ranger, J. Hulcr, J. Galko, and P. Kaňuch. 2020. Origin of non-native Xylosandrus germanus, an invasive pest ambrosia beetle in Europe and North America. Journal of Pest Science 94:553–562.

El Mokhefi, M., C. Kerdelhue, C. Burban, A. Battisti, G. Chakali, and M. Simonato. 2016. Genetic differentiation of the pine processionary moth at the southern edge of its range: contrasting patterns between mitochondrial and nuclear markers. Ecology and Evolution 6:4274–4288.

Estoup, A., and T. Guillemaud. 2010. Reconstructing routes of invasion using genetic data: why, how and so what? Molecular Ecology 19:4113–4130.

Etter, P. D., S. Bassham, P. A. Hohenlohe, E. A. Johnson, and W. A. Cresko. 2011. SNP discovery and genotyping for evolutionary genetics using RAD sequencing. Methods in Molecular Biology 772:157–178.

Folmer, O., M. Black, W. Hoeh, R. Lutz, and R. Vrijenhoek. 1994. DNA primers for amplification of mitochondrial cytochrome c oxidase subunit I from diverse metazoan invertebrates. Molecular Marine Biology and Biotechnology 3:294–299.

Galili, T. 2015. dendextend: an R package for visualizing, adjusting and comparing trees of hierarchical clustering. Bioinformatics 31:3718–3720.

Gallego, D., J. L. Lencina, H. Mas, J. Cevero, and M. Faccoli. 2017. First record of the granulate ambrosia beetle, Xylosandrus crassiusculus (Coleoptera: Curculionidae, Scolytinae), in the Iberian Peninsula. Zootaxa 4273:431–434.

Godefroid, M., A. Cruaud, J. P. Rossi, and J. Y. Rasplus. 2015. Assessing the risk of invasion by Tephritid fruit flies: intraspecific divergence matters. PLoS ONE 10:e0135209.

Guangchuang, Y. 2020. scatterpie: Scatter Pie Plot. R package version 0.1.5.

Guisan, A., W. Thuiller, and N. E. Zimmermann. 2017. Habitat suitability and distribution models with applications in R. Cambridge University Press.

Hagedorn, D. E. Z. 1908. Xyleborus mascarenus. Deutsche Ent. Zeitschr 379.

Hughes, M. A., J. J. Riggins, F. H. Koch, A. I. Cognato, C. Anderson, J. P. Formby, T. J. Dreaden, R. C. Ploetz, and J. A. Smith. 2017. No rest for the laurels: symbiotic invaders cause unprecedented damage to southern USA forests. Biological Invasions 19:2143–2157.

Hulcr, J., and L. L. Stelinski. 2017. The ambrosia symbiosis: from evolutionary ecology to practical management. Annual Review of Entomology 62:285–303.

Ito, M., and H. Kajimura. 2009. Phylogeography of an ambrosia beetle, Xylosandrus crassiusculus (Motschulsky) (Coleoptera: Curculionidae: Scolytinae), in Japan. Applied Entomology and Zoology 44:549–559.

Javal, M., E. Lombaert, T. Tsykun, C. Courtin, C. Kerdelhue, S. Prospero, A. Roques, and G. Roux. 2019. Deciphering the worldwide invasion of the Asian long-horned beetle: A recurrent invasion process from the native area together with a bridgehead effect. Molecular Ecology 28:951–967.

Jones, B. A., and S. M. McDermott. 2017. Health impacts of invasive species through an altered natural environment: assessing air pollution sinks as a causal pathway. Environmental and Resource Economics 71:23–43.

Kajtoch, Ł., M. Gronowska, R. Plewa, M. Kadej, A. Smolis, T. Jaworski, and J. M. Gutowski. 2022. A review of saproxylic beetle intra- and interspecific genetics: current state of the knowledge and perspectives. The European Zoological Journal 89:481–501.

Kavčič, A. 2018. First record of the Asian ambrosia beetle, Xylosandrus crassiusculus (Motschulsky) (Coleoptera: Curculionidae, Scolytinae), in Slovenia. Zootaxa 4483:191–193.

Kavčič, A., and M. de Groot. 2017. Pest risk analysis for the Asian Ambrosia Beetle (Xylosandrus crassiusculus (Motschulsky, 1866)). Slovenian Forestry Institute.

Kenis, M., M.-A. Auger-Rozenberg, A. Roques, L. Timms, C. Péré, M. J. W. Cock, J. Settele, S. Augustin, and C. Lopez-Vaamonde. 2008. Ecological effects of invasive alien insects. Biological Invasions 11:21–45.

Kirkendall, L. R. 2018. Invasive bark beetles (Coleoptera, Curculionidae, Scolytinae) in Chile and Argentina, including two species new for South America, and the correct identity of the Orthotomicus species in Chile and Argentina. Diversity 10.

Kumar, S., G. Stecher, M. Li, C. Knyaz, and K. Tamura. 2018. MEGA X: Molecular Evolutionary Genetics Analysis across Computing Platforms. Molecular Biology and Evolution 35:1547–1549.

LaBonte, J. R. 2010. Eradictation of an exotic ambrosia beetle, Xylosandrus crassiusculus (Motschulsky), in Oregon. Pages 41–43 in USDA Research Forum on Invasive Species.

Landi, L., D. Gómez, C. L. Braccini, V. A. Pereyra, S. M. Smith, and A. E. Marvaldi. 2017. Morphological and molecular identification of the invasive Xylosandrus crassiusculus (Coleoptera: Curculionidae: Scolytinae) and its South American range extending into Argentina and Uruguay. Annals of the Entomological Society of America 110:344–349.

Lesieur, V., E. Lombaert, T. Guillemaud, B. Courtial, W. Strong, A. Roques, and M. A. Auger-Rozenberg. 2018. The rapid spread of Leptoglossus occidentalis in Europe: a bridgehead invasion. Journal of Pest Science 92:189–200.

Li, H., and R. Durbin. 2009. Fast and accurate short read alignment with Burrows-Wheeler transform. Bioinformatics 25:1754–1760.

Lombaert, E., T. Guillemaud, J. M. Cornuet, T. Malausa, B. Facon, and A. Estoup. 2010. Bridgehead effect in the worldwide invasion of the biocontrol harlequin ladybird. PLoS ONE 5:e9743.

Nei, M. 1972. Genetic distance between populations. The American Naturalist 106:283–292.

Nel, W. J., Z. W. De Beer, M. J. Wingfield, and T. A. Duong. 2020. The granulate ambrosia beetle, Xylosandrus crassiusculus (Coleoptera: Curculionidae, Scolytinae), and its fungal symbiont found in South Africa. Zootaxa 4838:zootaxa 4838 4833 4837.

Paini, D. R., A. W. Sheppard, D. C. Cook, P. J. De Barro, S. P. Worner, and M. B. Thomas. 2016. Global threat to agriculture from invasive species. Proceedings of the National Academy of Science of the United States of America 113:7575–7579.

Paradis, E. 2010. pegas: an R package for population genetics with an integrated-modular approach. R package version 0.14. Bioinformatics 26:419–420.

Pattengale, N. D., M. Alipour, O. R. Bininda-Emonds, B. M. Moret, and A. Stamatakis. 2010. How many bootstrap replicates are necessary? Journal of Computational Biology 17:337–354.

Peer, K., and M. Taborsky. 2005. Outbreeding depression, but no inbreeding depression in haplodiploid ambrosia beetles with regular sibling mating. Evolution 59:317–323.

Pembleton, L. W., N. O. Cogan, and J. W. Forster. 2013. StAMPP: an R package for calculation of genetic differentiation and structure of mixed-ploidy level populations. Molecular Ecology Resources 13:946–952.

Pennachio, F., P. F. Roversi, V. Francardi, and E. Gatti. 2003. Xylosandrus crassiusculus (Motschulsky) a bark beetle new to Europe (Coleoptera Scolytidae). Redia 86:77–80.

R Core Team. 2018. R: a language and environment for statistical computing. R Foundation for Statistical Computing, Vienna, Austria.

Ramage, T., P. Martins-Simoes, G. Mialdea, R. Allemand, A. Duplouy, P. Rousse, N. Davies, G. K. Roderick, and S. Charlat. 2017. A DNA barcode-based survey of terrestrial arthropods in the Society Islands of French Polynesia: host diversity within the SymbioCode Project. European Journal of Taxonomy 272:1–13.

Ranger, C. M., M. E. Reding, P. B. Schultz, J. B. Oliver, S. D. Frank, K. M. Addesso, J. Hong Chong, B. Sampson, C. Werle, S. Gill, and C. Krause. 2016. Biology, ecology, and management of nonnative ambrosia beetles (Coleoptera: Curculionidae: Scolytinae) in ornamental plant nurseries. Journal of Integrated Pest Management 7:1–23.

Regupathy, A., and R. Ayyasamy. 2014. Occurrence of ambrosia beetles, Xylosandrus compactus (Eichh) and Xylosandrus crassiusculus (Motschulsky) on avocado in Tamil Nadu India: pest risk assessment. in Pacific Northwest Insect Management Conference, Portland, Oregon, USA.

Rochette, N. C., A. G. Rivera-Colon, and J. M. Catchen. 2019. Stacks 2: analytical methods for paired-end sequencing improve RADseq-based population genomics. Molecular Ecology 28:4737–4754.

Ronquist, F., M. Teslenko, P. van der Mark, D. L. Ayres, A. Darling, S. Hohna, B. Larget, L. Liu, M. A. Suchard, and J. P. Huelsenbeck. 2012. MrBayes 3.2: efficient Bayesian phylogenetic inference and model choice across a large model space. Systematic Biology 61:539–542.

Roques, A., R. Bellanger, J. B. Daubrée, C. Ducatillon, T. Urvois, and M.-A. Auger-Rozenberg. 2019. Les scolytes exotiques : une menace pour le maquis. Phytoma 727:16–20.

Rugman-Jones, P. F., M. Au, V. Ebrahimi, A. Eskalen, C. Gillett, D. Honsberger, D. Husein, M. G. Wright, F. Yousuf, and R. Stouthamer. 2020. One becomes two: second species of the Euwallacea fornicatus (Coleoptera: Curculionidae: Scolytinae) species complex is established on two Hawaiian islands. PeerJ 8:e9987.

Samuelson, G. A. 1981. A synopsis of Hawaiian Xyleborini (Coleoptera: Scolytidae). Pacific Insects 23:50–92.

Sardain, A., E. Sardain, and B. Leung. 2019. Global forecasts of shipping traffic and biological invasions to 2050. Nature Sustainability 2:274–282.

Schedl, K. E. 1953. Fauna Madagascariensis - III. Pages 67–106 Mémoires de l’Institut Scientifique de Madagascar - Série E - Tome III.

Schrieber, K., and S. Lachmuth. 2017. The genetic paradox of invasions revisited: the potential role of inbreeding x environment interactions in invasion success. Biological Reviews of the Cambridge Philosophical Society 92:939–952.

Seebens, H., T. M. Blackburn, E. E. Dyer, P. Genovesi, P. E. Hulme, J. M. Jeschke, S. Pagad, P. Pyšek, M. Winter, M. Arianoutsou, S. Bacher, B. Blasius, G. Brundu, C. Capinha, L. Celesti-Grapow, W. Dawson, S. Dullinger, N. Fuentes, H. Jager, J. Kartesz, M. Kenis, H. Kreft, I. Kuhn, B. Lenzner, A. Liebhold, A. Mosena, D. Moser, M. Nishino, D. Pearman, J. Pergl, W. Rabitsch, J. Rojas-Sandoval, A. Roques, S. Rorke, S. Rossinelli, H. E. Roy, R. Scalera, S. Schindler, K. Stajerova, B. Tokarska-Guzik, M. van Kleunen, K. Walker, P. Weigelt, T. Yamanaka, and F. Essl. 2017. No saturation in the accumulation of alien species worldwide. Nature Communications 8:14435.

Simberloff, D., J. L. Martin, P. Genovesi, V. Maris, D. A. Wardle, J. Aronson, F. Courchamp, B. Galil, E. Garcia-Berthou, M. Pascal, P. Pyšek, R. Sousa, E. Tabacchi, and M. Vila. 2013. Impacts of biological invasions: what’s what and the way forward. Trends in Ecology and Evolution 28:58–66.

Sire, L., D. Gey, R. Debruyne, T. Noblecourt, F. Soldati, T. Barnouin, G. Parmain, C. Bouget, C. Lopez-Vaamonde, and R. Rougerie. 2019. The challenge of DNA barcoding saproxylic beetles in natural history collections - Exploring the potential of parallel multiplex sequencing with Illumina MiSeq. Frontiers in Ecology and Evolution 7.

Sloan, D. B., J. C. Havird, and J. Sharbrough. 2017. The on-again, off-again relationship between mitochondrial genomes and species boundaries. Molecular Ecology 26:2212–2236.

Smith, S. M., and A. I. Cognato. 2022. New non-native pseudocryptic Cyclorhipidion species (Coleoptera: Curculionidae: Scolytinae: Xyleborini) found in the United States as revealed in a multigene phylogeny. Insect Systematics and Diversity 6.

Smith, S. M., D. F. Gomez, R. A. Beaver, J. Hulcr, and A. I. Cognato. 2019. Reassessment of the species in the Euwallacea fornicatus (Coleoptera: Curculionidae: Scolytinae) complex after the rediscovery of the “lost” type specimen. Insects 10.

Stamatakis, A. 2014. RAxML version 8: a tool for phylogenetic analysis and post-analysis of large phylogenies. Bioinformatics 30:1312–1313.

Storer, C., A. Payton, S. McDaniel, B. Jordal, and J. Hulcr. 2017. Cryptic genetic variation in an inbreeding and cosmopolitan pest, Xylosandrus crassiusculus, revealed using ddRADseq. Ecology and Evolution 7:10974–10986.

Stouthamer, R., P. Rugman-Jones, P. Q. Thu, A. Eskalen, T. Thibault, J. Hulcr, L.-J. Wang, B. H. Jordal, C.-Y. Chen, M. Cooperband, C.-S. Lin, N. Kamata, S.-S. Lu, H. Masuya, Z. Mendel, R. Rabaglia, S. Sanguansub, H.-H. Shih, W. Sittichaya, and S. Zong. 2017. Tracing the origin of a cryptic invader: phylogeography of the Euwallacea fornicatus (Coleoptera: Curculionidae: Scolytinae) species complex. Agricultural and Forest Entomology 19:366–375.

Tang, J., K. Mao, H. Zhang, X. Xu, X. Xu, H. Guo, and B. Li. 2022. Multiple introductions and genetic admixture facilitate the successful invasion of Plantago virginica into China. Biological Invasions.

Toews, D. P. L., and A. Brelsford. 2012. The biogeography of mitochondrial and nuclear discordance in animals. Molecular Ecology 21:3907–3930.

UNCTAD. 2020. Review of maritime transport 2020. United Nations, Geneva.

Urvois, T., C. Perrier, A. Roques, L. Sauné, C. Courtin, Y. Li, A. J. Johnson, J. Hulcr, M.-A. Auger-Rozenberg, and C. Kerdelhué. 2022. A first inference of the phylogeography of the worldwide invader Xylosandrus compactus Journal of Pest Science 95:1217–1231.

Vanderpool, D., R. R. Bracewell, and J. P. McCutcheon. 2018. Know your farmer: ancient origins and multiple independent domestications of ambrosia beetle fungal cultivars. Molecular Ecology 27:2077–2094.

Weir, B. S., and C. C. Cockerham. 1984. Estimating F-statistics for the analysis of population structure. Evolution 38:1358–1370.

Wickham, H. 2016. ggplot2: Elegant graphics for data analysis. Springer-Verlag New York.

Wright, S. 1951. The genetical structure of populations. Annals of Eugenics 15:323–354.

Zheng, X. 2013. Statistical prediction of HLA alleles and relatedness analysis in genome-wide association studies. University of Washington, plWashington.

Zheng, X., D. Levine, J. Shen, S. M. Gogarten, C. Laurie, and B. S. Weir. 2012. A high-performance computing toolset for relatedness and principal component analysis of SNP data. Bioinformatics 28:3326–3328.

