## Supplementary Figures for "The worldwide invasion history of a pest ambrosia beetle inferred using population genomics"

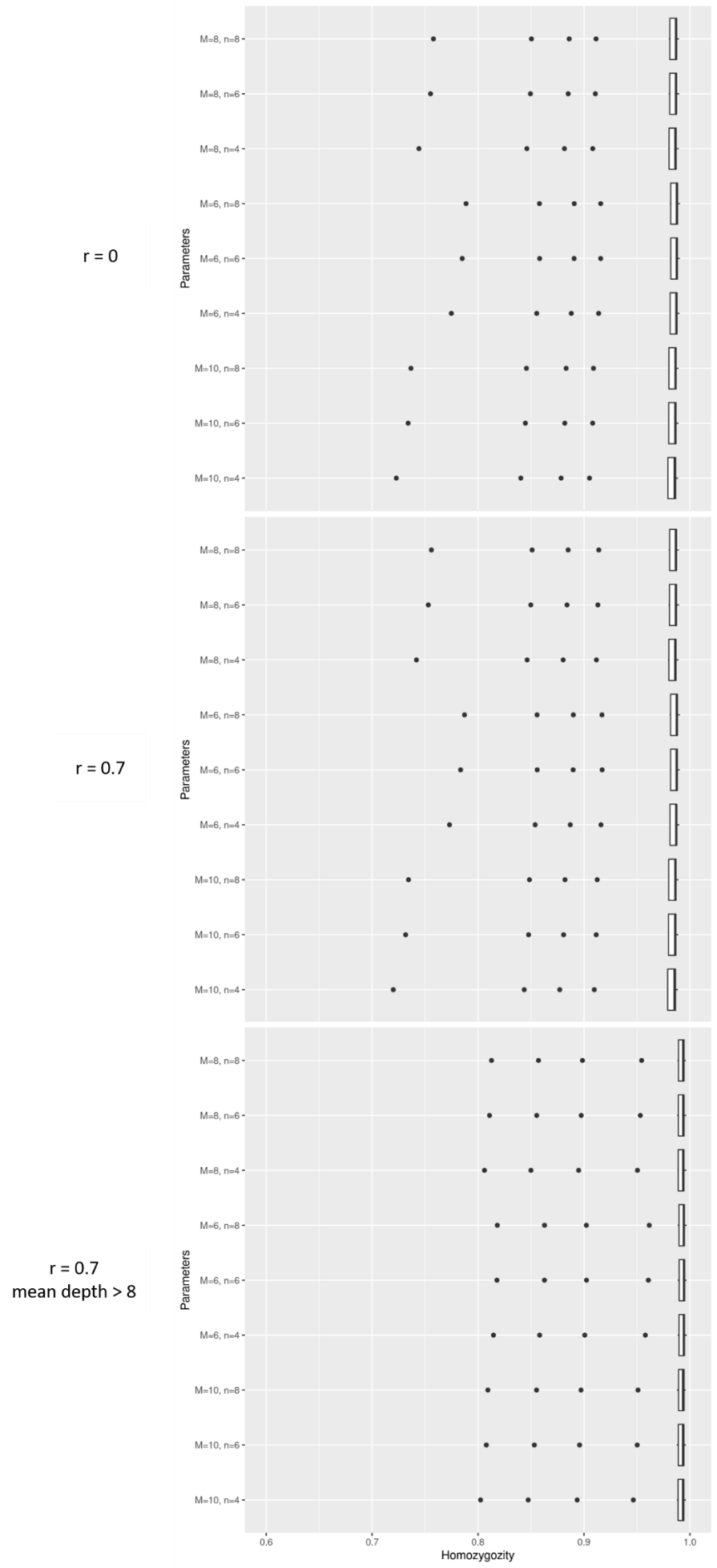

Supplementary Figure 1: Boxplots of the individual homozygosity rate for the different M and n combination tested (M = 6, 8 and 10, n = 4, 6 and 8) for three filtering conditions (without filtering, r = 0.7 and r = 0.7 and excluding loci with a mean depth lower than 8).

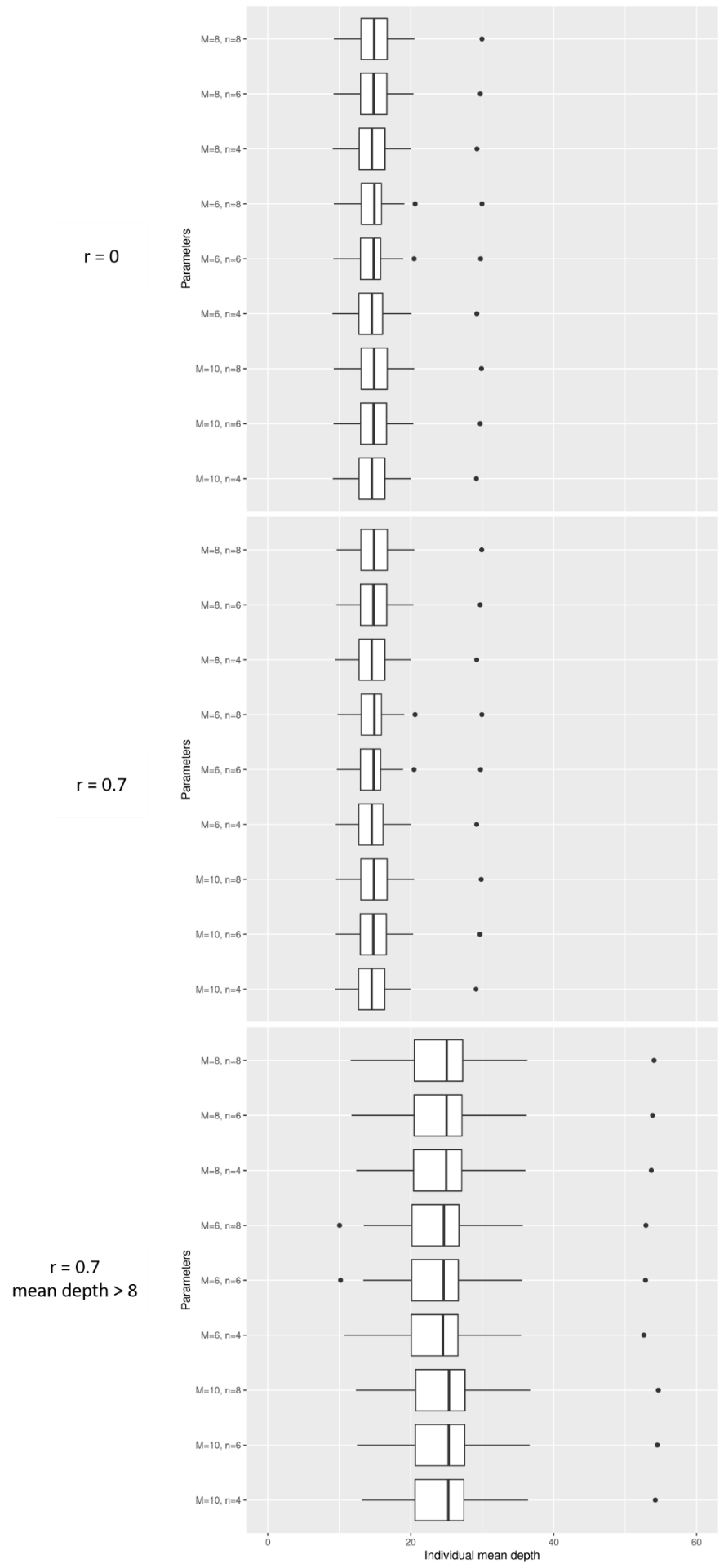

Supplementary Figure 2: Boxplots of the individual mean depth for the different M and n combination tested (M = 6, 8 and 10, n = 4, 6 and 8) for three filtering conditions (without filtering, r = 0.7 and r = 0.7 and excluding loci with a mean depth lower than 8).

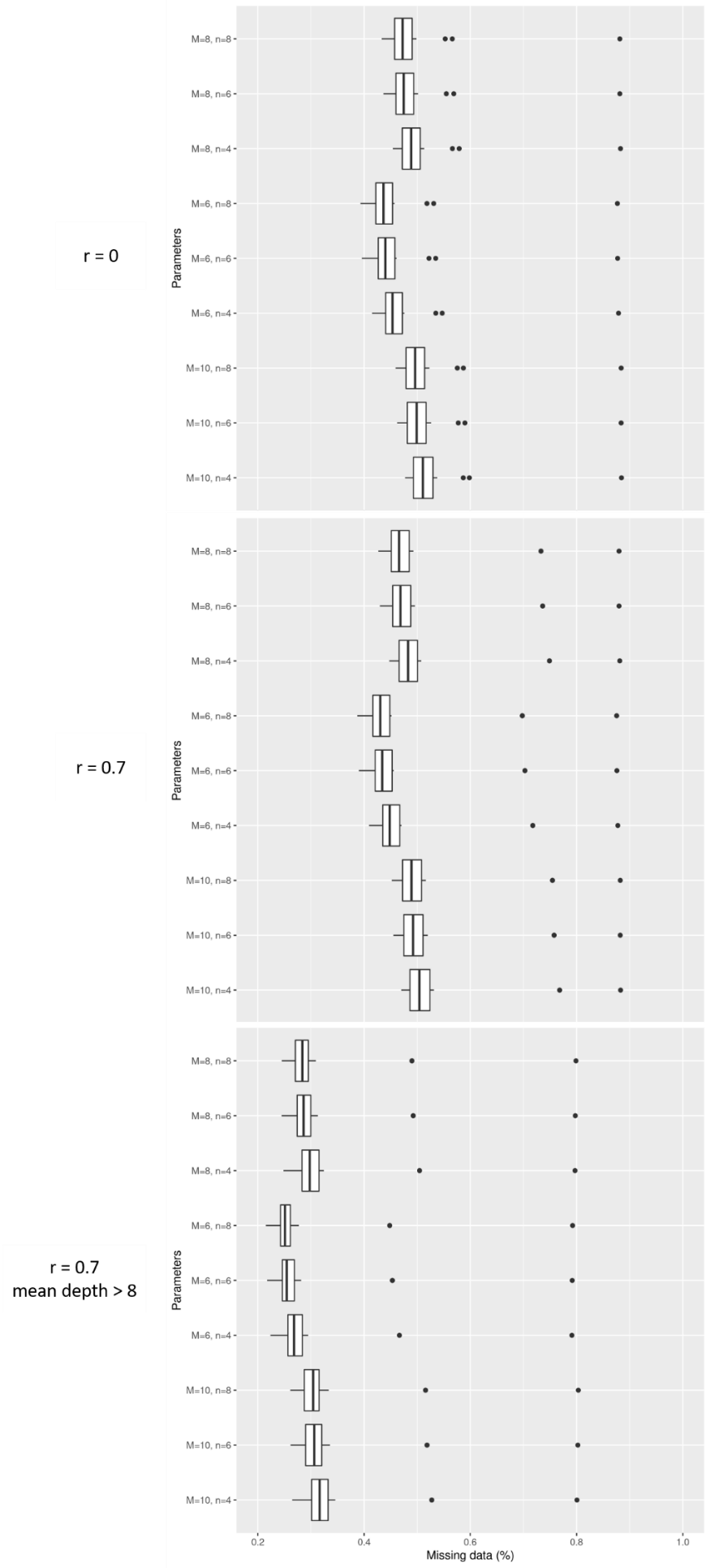

Supplementary Figure 3: Boxplots of the individual missing data (%) for the different M and n combination tested (M = 6, 8 and 10, n = 4, 6 and 8) for three filtering conditions (without filtering, r = 0.7 and r = 0.7 and excluding loci with a mean depth lower than 8).

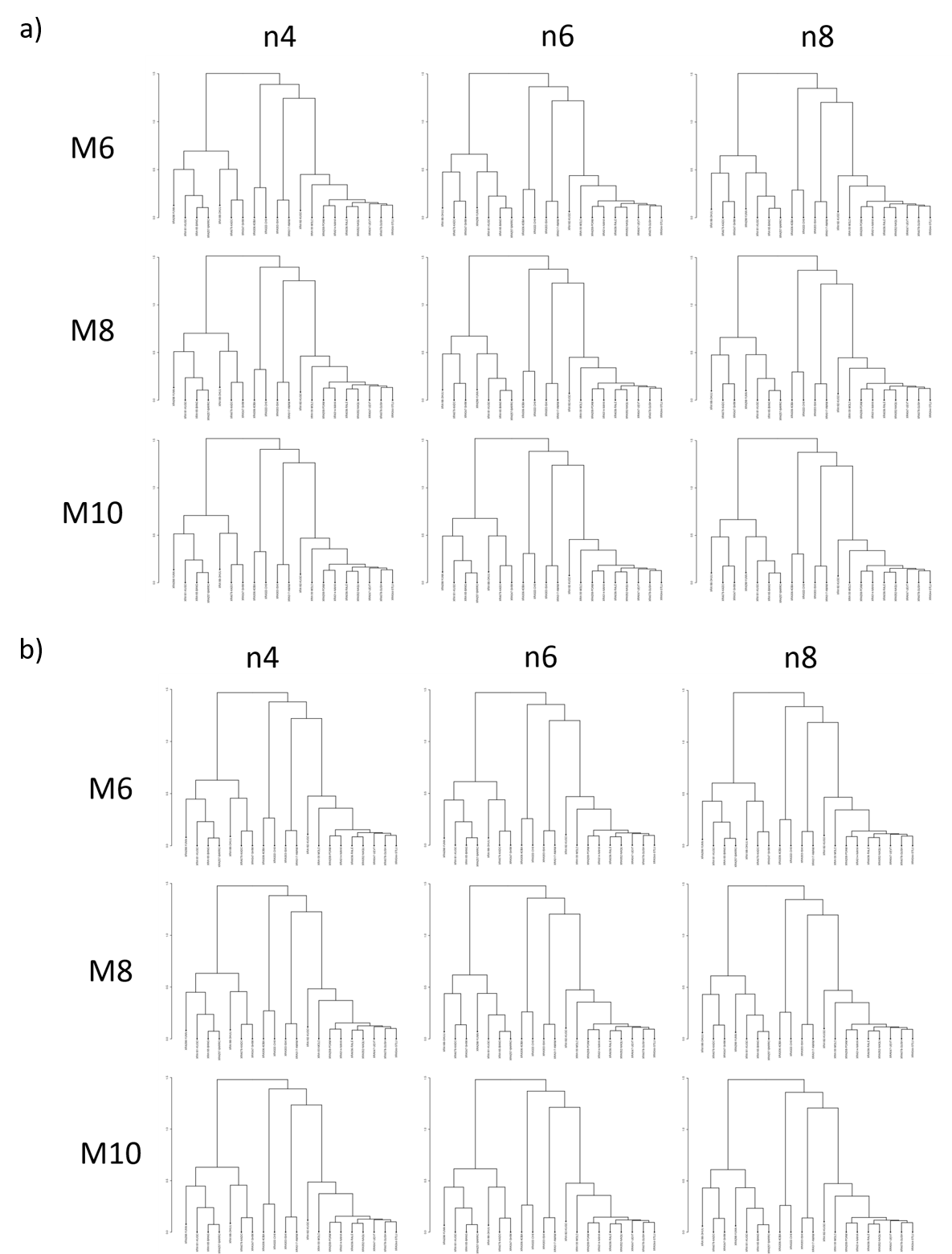

Supplementary Figure 4: Dendrograms obtained in the pre-analysis for the different M and n combination for a) r = 0 and without filtering on depth and b) r = 0.7 and after excluding loci with a mean depth lower than 8.

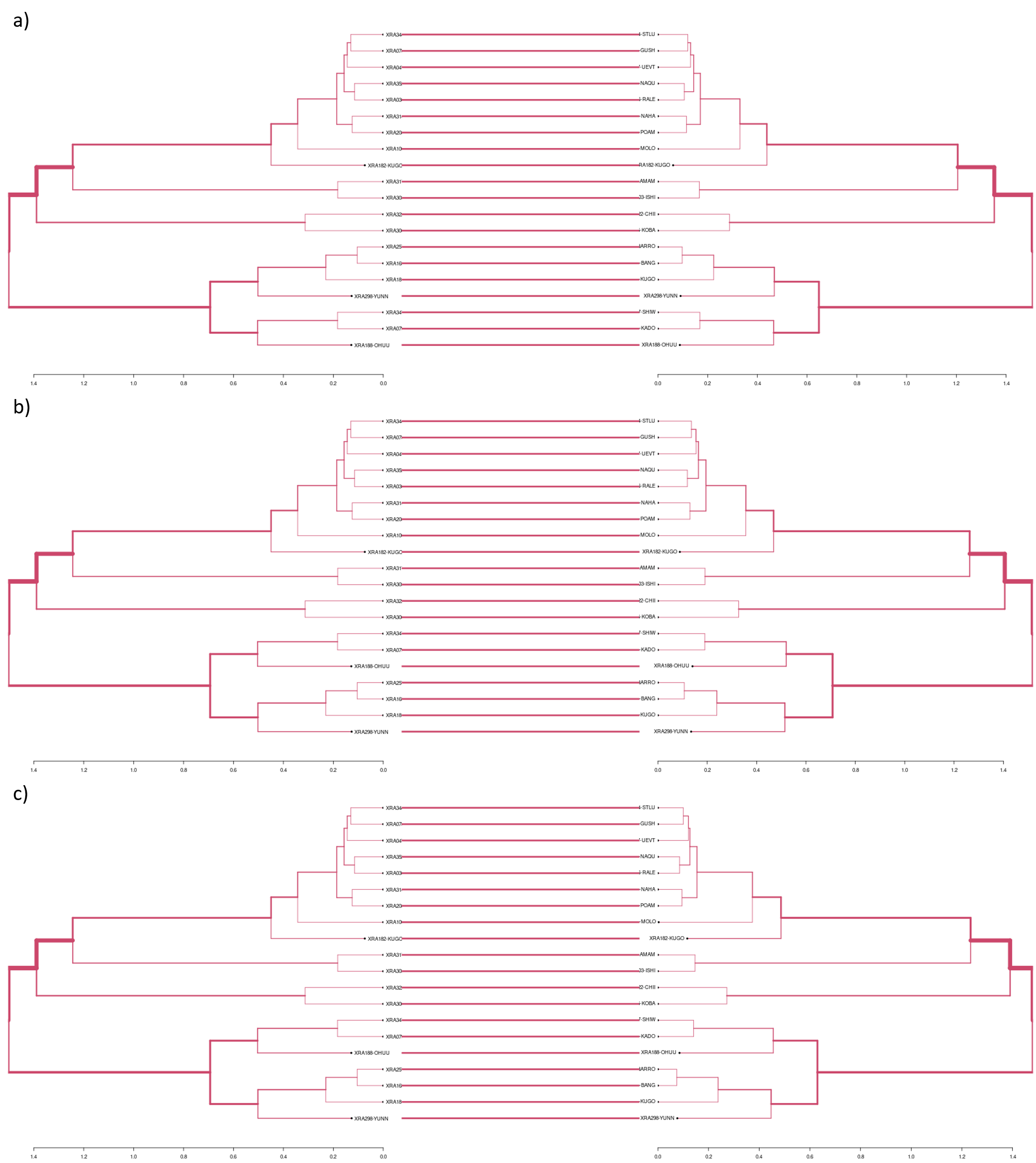

Supplementary Figure 5: Tanglegram plots comparing the dendrograms obtained in the pre-analysis with different parameter combinations: a) M = 6, n = 4, r = 0 and M = 6, n = 8, r = 0, b) M = 6, n = 4, r = 0 and M = 8, n = 4, r = 0 and c) M = 6, n = 4, r = 0 and M = 6, n = 4, r = 0.7, excluding the loci with a mean depth lower than 8. The entanglement value for each figure is 0, showing no difference between all dendrograms.

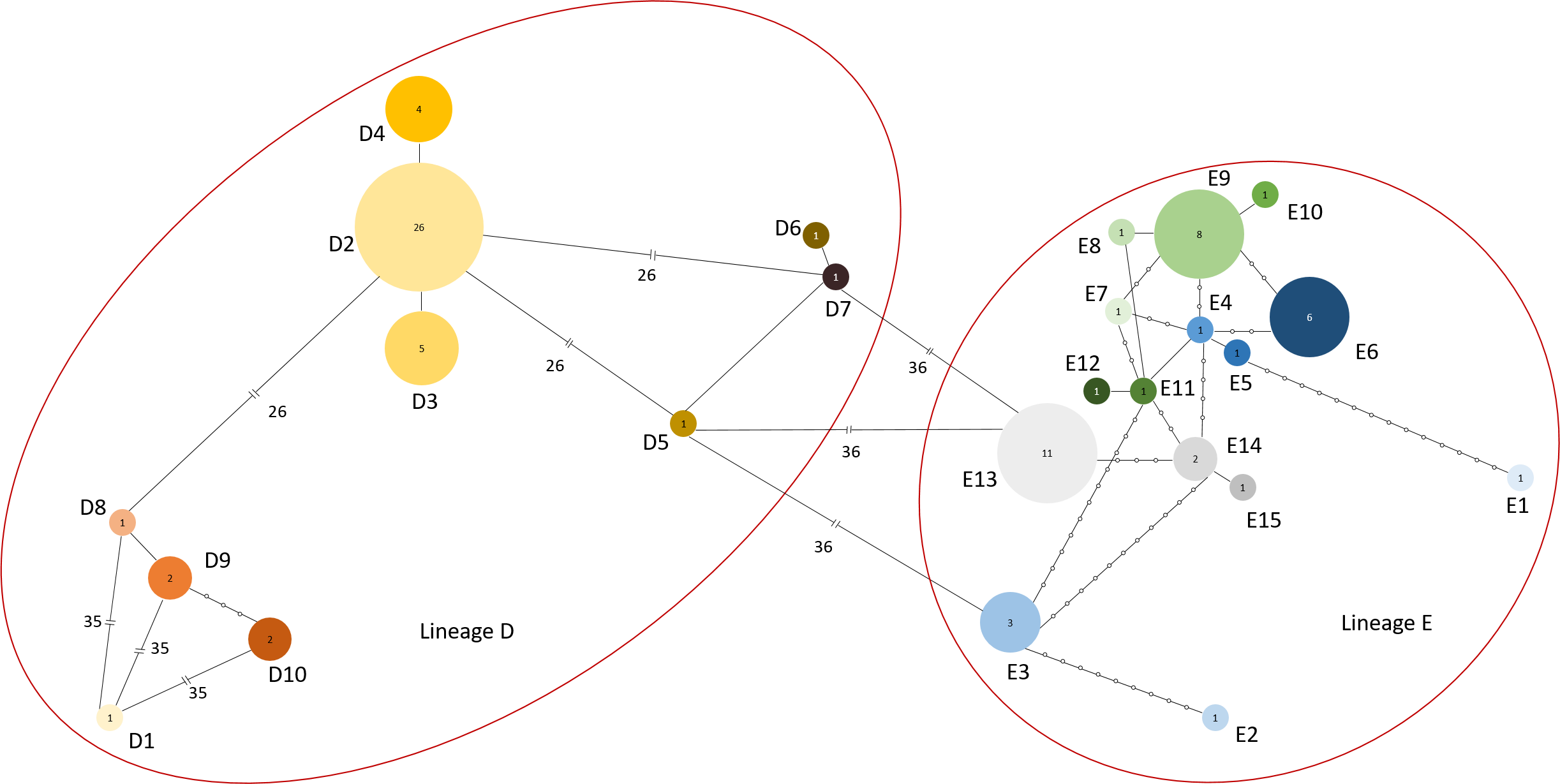

Supplementary Figure 6: Median-joining network representing Xylosandrus crassiusculus’ mitochondrial cluster 1 based on COI sequences.

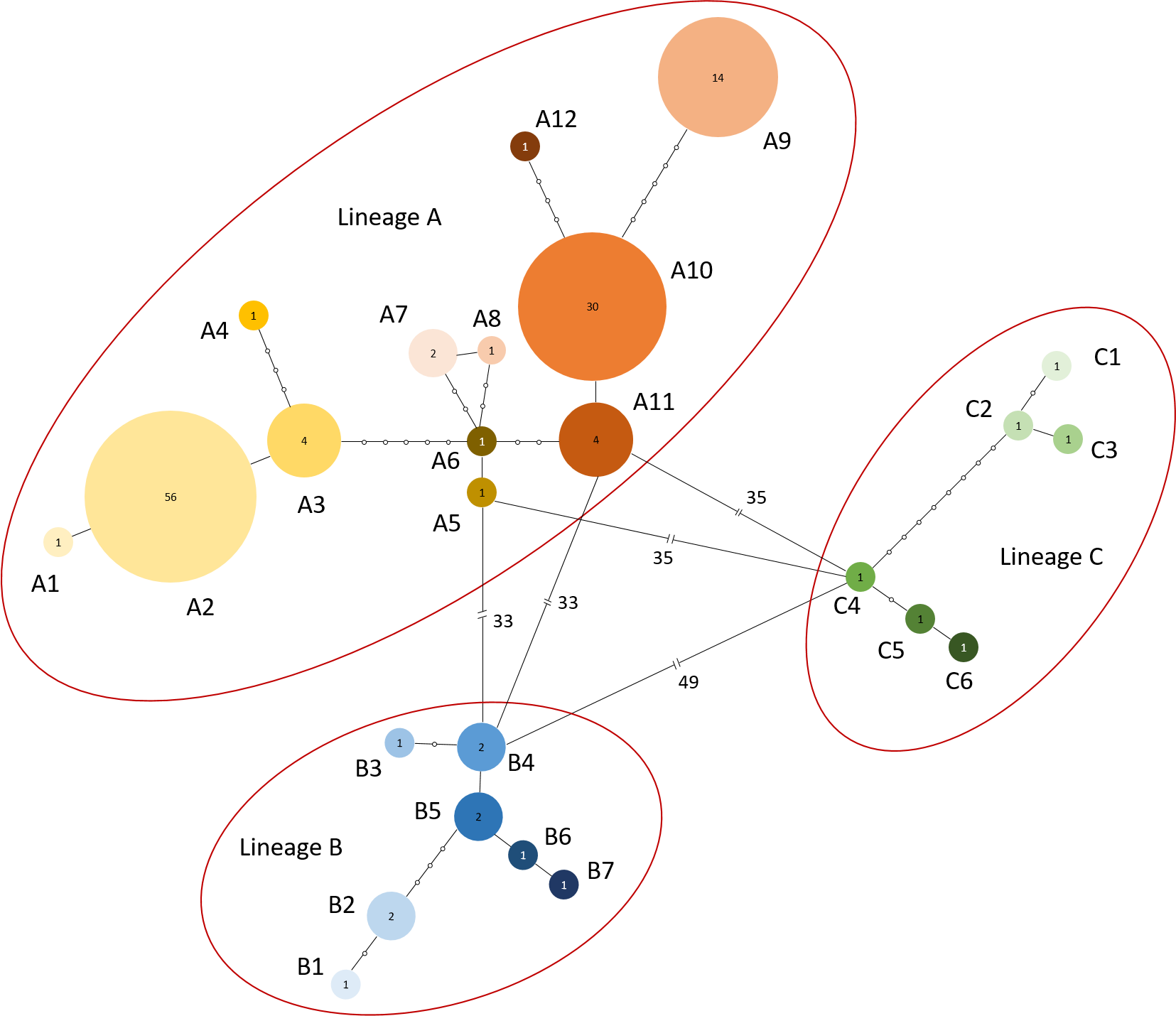

Supplementary Figure 7: Median-joining network representing Xylosandrus crassiusculus’ mitochondrial cluster 2 based on COI sequences.

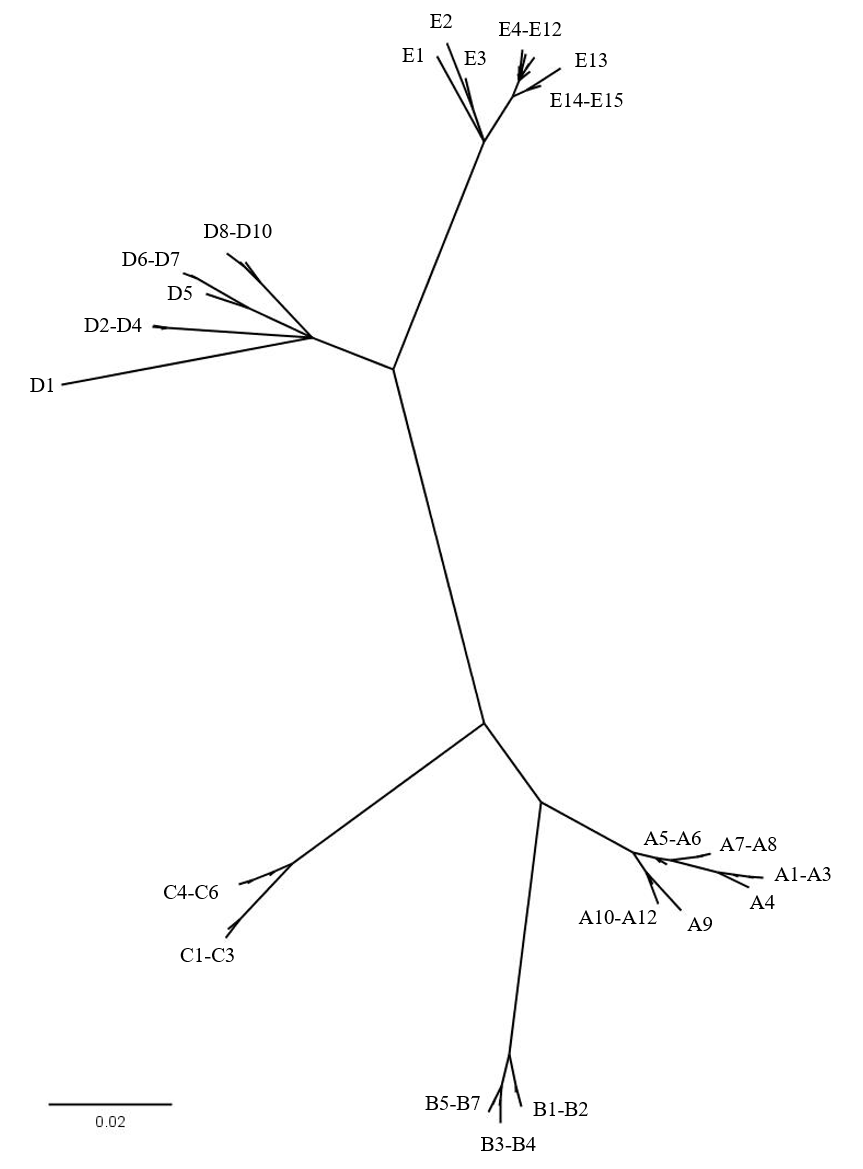

Supplementary Figure 8: Radial Bayesian tree based on X. crassiusculus’ COI sequences built with MrBayes 3.2.7 without outgroup.

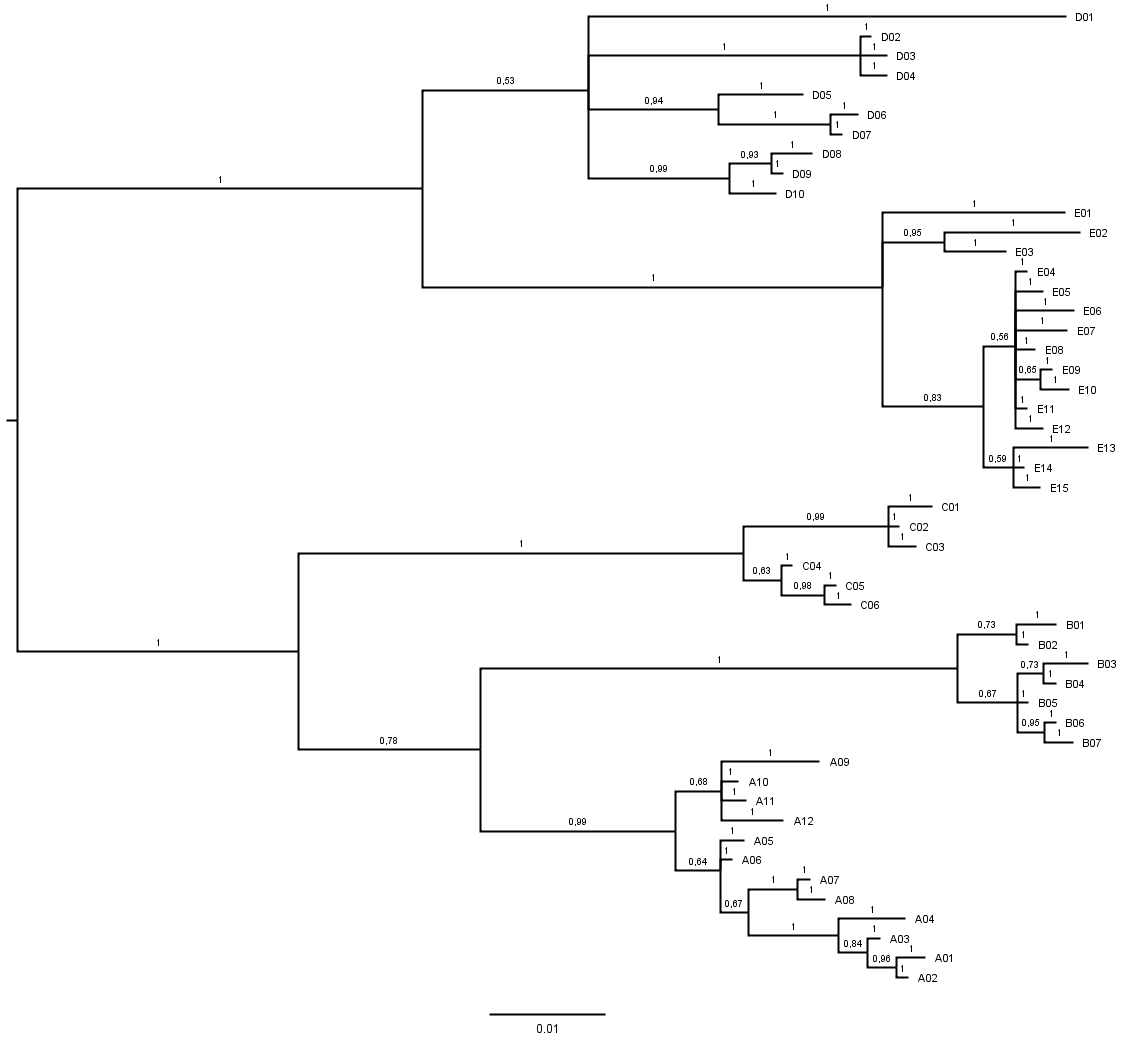

Supplementary Figure 9: Bayesian tree based on X. crassiusculus’ COI sequences built with MrBayes 3.2.7 without outgroup.

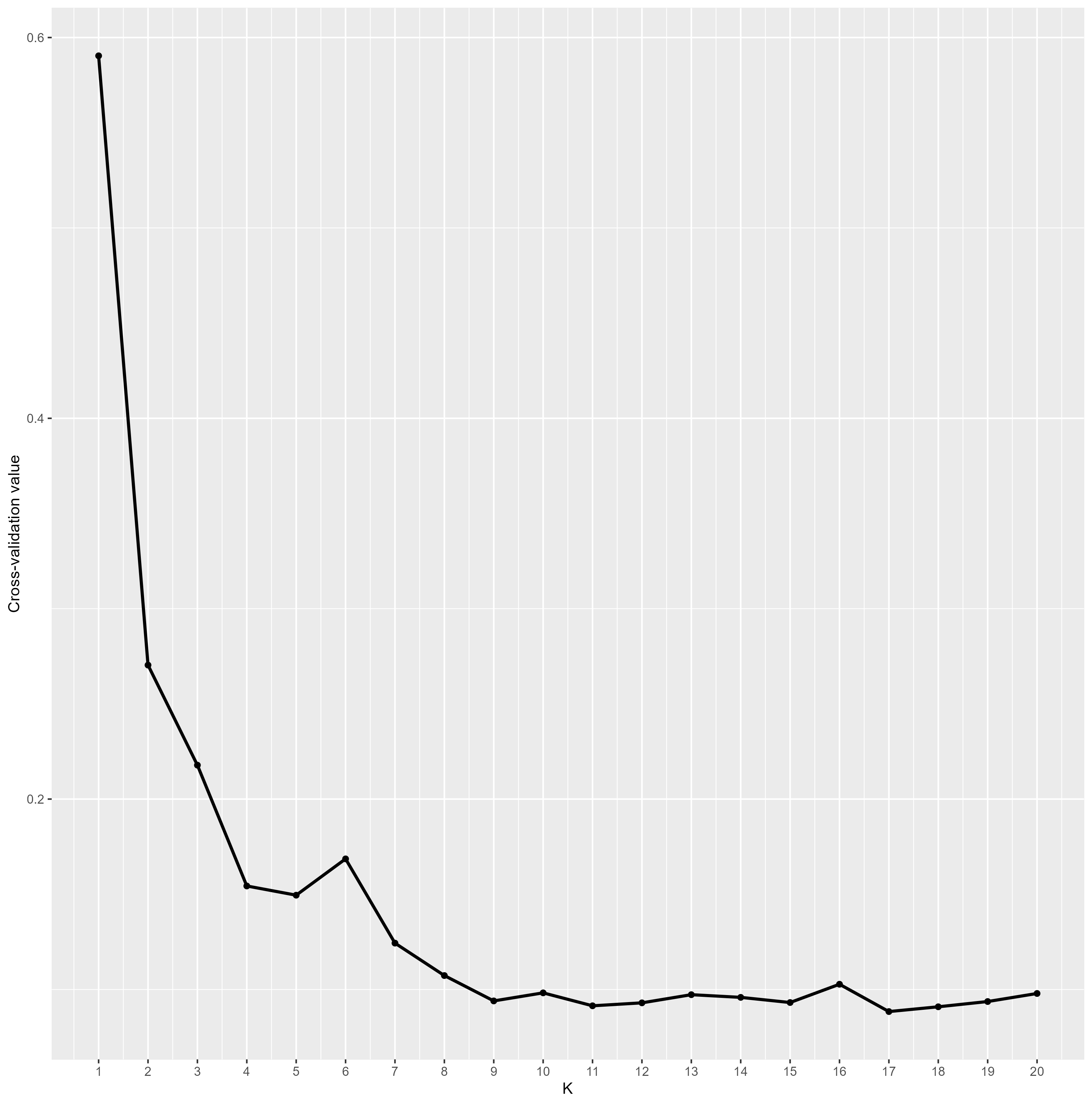
 Supplementary Figure 10: Cross-validation plot for K ranging from 1 to 20.
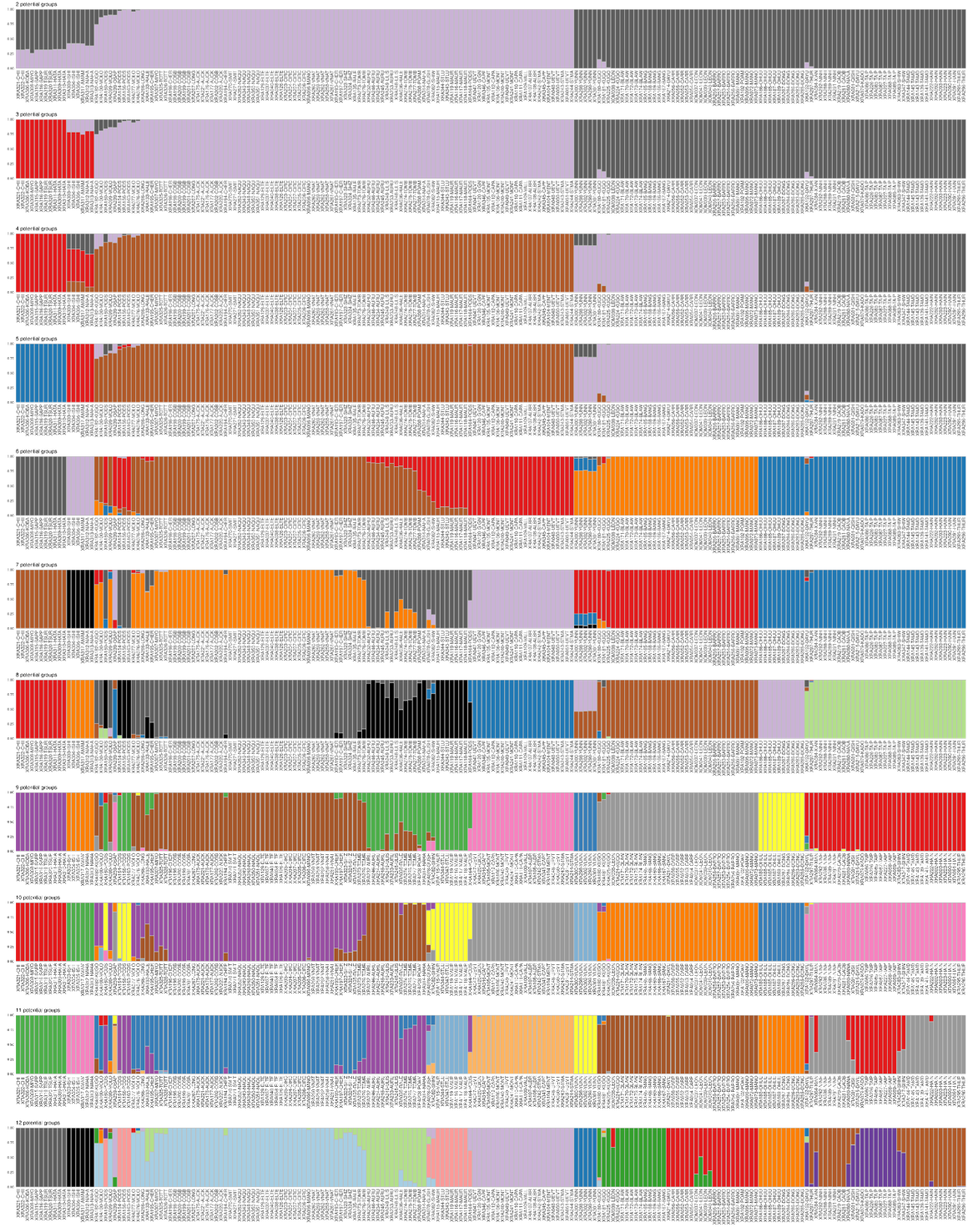
 Supplementary Figure 11: Admixture plots for K ranging from 2 to 12. Specimens are ordered according to their order in the clustering tree in Figure 8.

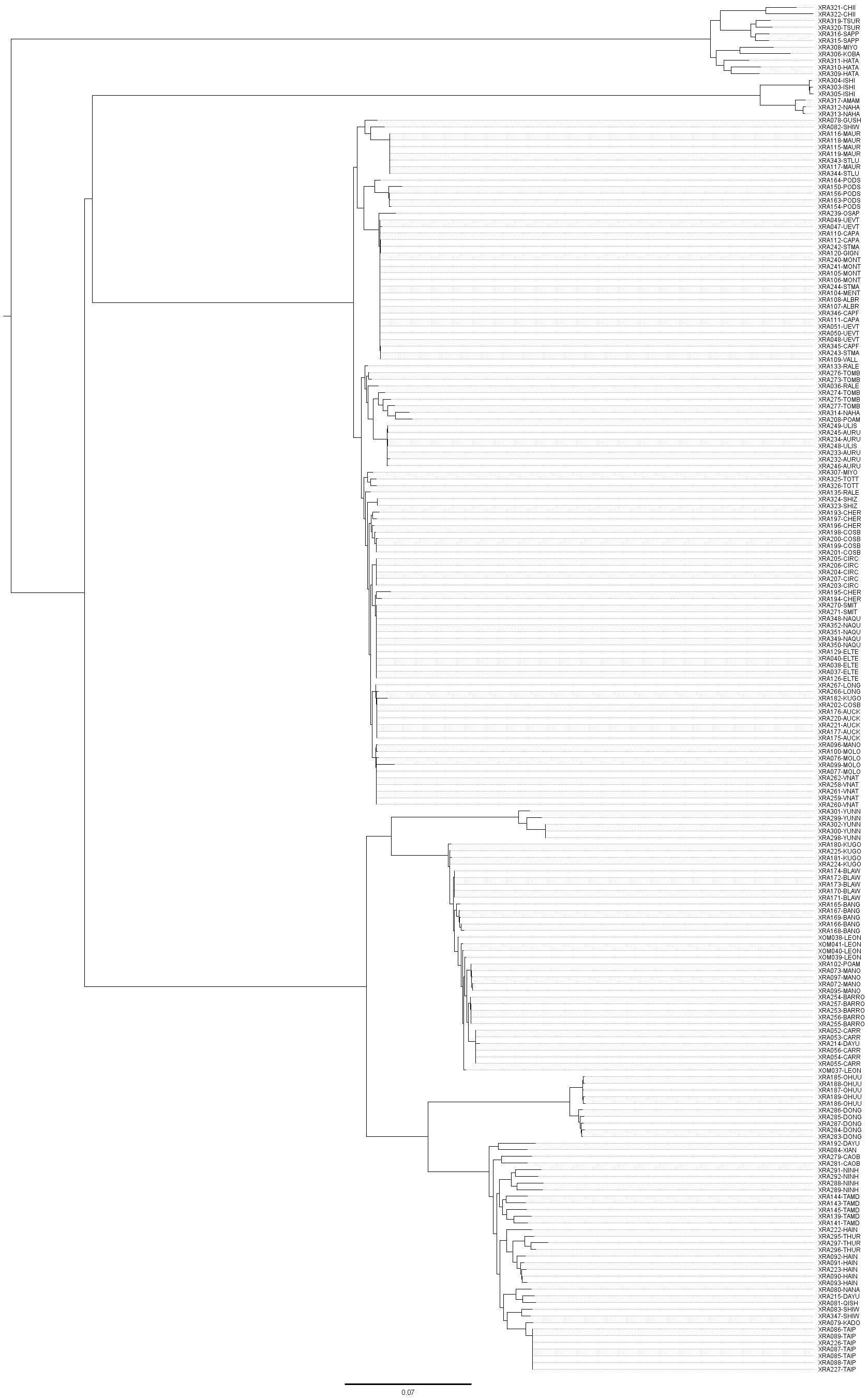

Supplementary Figure 12: Maximum likelihood tree based on X. crassiusculus’ RAD sequencing data performed with RAxML 8.2.1.
